# Human cellular homeostasis buffers *trans*-acting translational effects of heterologous gene expression with very different codon usage bias

**DOI:** 10.1101/2021.12.09.471957

**Authors:** Arthur J Jallet, Antonin Demange, Fiona Leblay, Mathilde Decourcelle, Khadija El Koulali, Marion AL Picard, Ignacio G Bravo

## Abstract

The frequency of synonymous codons in protein coding genes is non-random and varies both between species and between genes within species. Whether this codon usage bias (CUBias) reflects underlying neutral mutational processes or is instead shaped by selection remains an open debate, especially regarding the role of selection for enhanced protein production. Variation in CUBias of a gene (be it natural synonymous mutations or biotechnological synonymous recoding) can have an enormous impact on its expression by diverse *cis*-acting mechanisms. But expression of genes with extreme CUBias can also lead to strong phenotypic effects by altering the overall intracellular translation homeostasis *via* competition for ribosomal machinery or tRNA depletion. In this study, we expressed at high levels in human cells six different synonymous versions of a gene and used matched transcriptomic and proteomic data to evaluate the impact of CUBias of the heterologous gene on the translation of cellular transcripts. Our experimental design focused specifically on differences during translation elongation. Response to expression of the different synonymous sequences was assessed by various approaches, ranging from analyses performed on a per-gene basis to more integrated approaches of the cell as a whole. We observe that the transcriptome displayed substantial changes as a result of heterologous gene expression by triggering an intense antiviral and inflammatory response, but that changes in the proteomes were very modest. Most importantly we notice that changes in translation efficiency of cellular transcripts were not associated with the direction of the CUBias of the heterologous sequences, thereby providing only limited support for *trans*-acting effects of synonymous changes. We interpret that, in human cells in culture, changes in CUBias can lead to important *cis*-acting effects in gene expression, but that cellular homeostasis can buffer the phenotypic impact of overexpression of heterologous genes with extreme CUBias.

## Introduction

The genetic code allows translating information stored in gene nucleotide sequences into protein sequences. This code is redundant because the standard twenty amino acids are encoded by 61 codons. Codons encoding the same amino acid are called synonymous codons and amino acids can be encoded by up to six synonymous codons. It was early noted that synonymous codons are not used at random but instead certain synonymous codons are over-represented in the genome compared to the null expectation of uniform use. This non-random usage of synonymous codons is referred to as codon usage bias (CUBias). CUBias widely varies across species (**Ikemura, 1982; Kanaya et al, 1999; Novoa et al, 2019**) as well as between genes within the same genome (**Gouy & Gautier, 1982; Sharp & Li, 1986; Duret, 2002**).

Two non-mutually exclusive hypotheses have been formulated to explain the existence of CUBias (**Hershberg & Petrov, 2008; Plotkin & Kudla, 2011**). The first one is the *mutational bias hypothesis*, which is usually conceived as a neutral explanation as it does not involve any selective process shaping CUBias. According to this view, the CUBias of a coding sequence simply arises because of local biases in the mutation spectrum during DNA replication and/or repair. Since such mutational biases are expected to be similar for coding and non-coding parts of the genome, CUBias should thus mirror the nucleotide composition of non-coding sequences nearby. This idea of CUBias being an epiphenomenon of mutational processes is supported by several lines of evidence. For example, GC composition of exons is usually similar to that of the introns in the same gene (**Chamary, Parmley & Hurst, 2006**) or to that of intergenic sequences in its vicinity (**Chen et al, 2004**), illustrating that mapping or not onto a translated region is not the major determinant of the nucleotide composition bias. At broader scale, evidence of such co-variation of nucleotide composition between coding and non-coding regions is obvious in the so-called isochores, present in the genomes of many vertebrates. Isochores are long chromosome stretches enriched in either AT or GC nucleotides (**Caspersson et al, 1968**). This strong compositional bias along large genomic regions is maintained over evolution, so that the physical mapping of a gene onto a given isochore is the most important determinant of nucleotide composition and therefore of CUBias (**Holmquist, 1989; Duret, 2002; Pouyet et al, 2017**). Finally, local CUBias in vertebrate genomes is further influenced by the distance to recombination hotspots and could reflect the intensity of the GC-biased gene conversion around the considered loci (**Pouyet et al, 2017**).

In the second hypothesis, known as *translational selection*, synonymous mutations are considered subject to selection as they can provide phenotypic variation and ultimately differences in fitness by means of variation in protein amount and/or quality during the gene expression process, mostly during translation. According to this view, selection can discriminate between synonymous codons either because of their different decoding speed, their different consequences on co-translational folding of the translated protein (an effect probably mediated by local decoding speed itself – **Pechmann & Fryman, 2013; Yu et al, 2015**) or the different accuracy for codon-anticodon pairing (**Stoletzki & Eyre-Walker, 2007; Drummond & Wilke, 2008; Drummond & Wilke, 2009; Walsh et al, 2020**). The proposed involvement of CUBias on translational efficiency is strongly supported by the fact that genes preferentially using the most frequent synonymous codons are usually those expressed at higher levels and decoded by more abundant transfer RNAs (tRNAs). This association between codon frequency and tRNA abundance has been extensively reported in multiple fast growing, unicellular organisms such as *E. coli* (**Ikemura, 1981; Sharp & Li, 1986; Tuller et al, 2010**) or *S. cerevisiae* (**Ikemura, 1982; Akashi, 2003**; **Tuller et al, 2010**), but also in multicellular organisms such as *C. elegans* (**Duret, 2000**) and *D. melanogaster* (**Duret & Mouchiroud, 1999**). This association supports that CUBias can be shaped by selection to optimize protein production. Technical advances, such as ribosome profiling, confirm that synonymous mutations play a role in how efficiently genes are translated, and that this differential efficiency is directly linked to tRNA availability (**Hia & Takeuchi, 2021**, but see **Charneski & Hurst, 2013**). In yeast, translation efficiency is higher when codon frequency matches tRNA availability (**Riba et al, 2019**), and decoding time of individual codons is anti-correlated with the availability of the corresponding cognate tRNA (**Gardin et al, 2014**; **Weinberg et al, 2016**). While supported in the above-cited model species, translational selection in mammalian genomes is a much more debated topic (**Urrutia & Hurst, 2001**) and if exists, its influence seems to be much weaker than that of mutational processes (**Kanaya et al, 2001; Vogel et al, 2010**). Further, the importance of GC-biased gene conversion and isochore compartmentation on shaping local genome composition is probably strong enough to blur most of the signatures left within coding sequences by selection (**Pouyet et al, 2017**).

The translational selection hypothesis proposes that the CUBias of a gene can modulate its own translation efficiency, an impact usually referred to as *cis*-effects. But it has been proposed that the CUBias of a gene can exert effects on the expression and translation efficiency of other genes, by means of *trans*-effects. Indeed, translation is the most expensive individual step of the gene expression process (**Lynch & Marinov, 2015**) and the availability of the complex and costly translational machinery is the overall limitation for protein synthesis. The different cellular transcripts face thus a direct competition for finite translational resources, such as ribosomes or tRNAs (**Li G et al, 2014**). As a result, there are large differences between transcripts in their ribosomal binding ability, so that not necessarily the most abundant mRNAs are those that are more effective at recruiting ribosomes and initiating translation, resulting in loss of translation opportunity for other mRNAs (**Callens et al, 2021**). Further, when abundant and actively ribosome-recruiting mRNA species perform poorly at translation elongation – because of a poor CUBias, for instance –, ribosomal pausing leads to accumulation of slowly proceeding ribosomes, amplifying ribosome sequestration and further reducing the pool of free ribosomes available for translation of other mRNAs (**Shah et al, 2013; Pelechano, Wei & Steinmetz, 2015**). From a practical perspective, competition between mRNAs to access shared resources such as tRNAs can be considered under a perspective of cellular economy, using a supply-and-demand reasoning. In this perspective, the pool of transcripts exerts a *demand* for being translated, requiring their codons to be decoded by the corresponding cognate anticodons, which represent the *offer* (**Gingold & Pilpel, 2011; Gingold, Dahan & Pilpel, 2012**).

Since *trans*-acting effects of gene overexpression can hamper translation of essential genes, it is conceivable that genes that require high expression levels have been selected to be encoded with a particular CUBias, to decrease the burden caused onto other genes arising from their high expression levels, or to avoid suffering from *trans* acting effects caused by high expression level of other genes. Two studies in *E. coli* strongly support this hypothesis of *trans-*acting effects mediated by competition between mRNAs: i) first, expressing different synonymous versions of the green fluorescent protein (GFP) did not result in major changes in its own translation *-i.e*., no *cis-* effects-, but instead led to significant *trans-*effects on global translation efficiency (**Kudla et al, 2009**); ii) second, the global negative effects associated with translating highly expressed genes that use rare codons can be reversed if the availability of the cognate tRNAs is increased to meet the demand (**Frumkin et al, 2018**). Results consistent with such *trans-* effects have been also communicated for yeast (**Pop et al, 2014**).

In this study we aim at understanding the extent and strength of *trans*-acting effects by analyzing the different cellular changes incurred as a function of heterologous expression of genes with different CUBias. We use human HEK293 cells as model system and combine transcriptomic and proteomic data to identify patterns consistent with *trans*-acting effects on translation efficiency mediated by competition for the translation machinery. The analyses of the *cis*-acting effects of codon recoding have been the focus of a separate study (**Picard et al, 2022**). Here we analyze both individual gene translation efficiency and overall translation efficiency at the whole-cell scale. Globally our results show that heterologous gene overexpression triggers changes of large extent in the cellular transcriptome. In contrast, changes in the cellular proteome are more discrete and do not show any evident global trend with regards to CUBias of the heterologous genes. More importantly, our results suggest that cellular homeostasis can largely buffer the translational effects of gene overexpression in our experimental system. This study thereby provides only limited support for the hypothesis of *trans*-acting effects arising from directional competition for limited translational resources, associated to CUBias in humans.

## Results

### Transfection and heterologous gene expression lead to substantial changes in the cellular transcriptome, independently of the codon usage bias of the heterologous genes

In the present study we have transfected human HEK293 cells with a pcDNA3.1 plasmid containing different synonymous versions of the *shble* gene (**Fig. 1A**) and analysed transcriptional and translational differential changes associated to heterologous gene expression. *Shble* sequences were translationally coupled to the *egfp* gene, under the control of a strong promoter. We performed three rounds of transfection experiments, each one including six *shble* synonymous versions, one transfection control with the Empty vector (*i.e*. our plasmid backbone with no *shble* sequence and encoding only for the *egfp and NeoR* genes, with identical sequences for all constructs), as well as one MOCK control with mock-transfected cells. Transcriptomic analyses were run in biological duplicates for each experiment, so that we obtained a total of 48 cellular transcriptomes, that were latter collapsed to 24 by averaging TPM values of duplicates (see Methods). For proteome analyses, the biological duplicates in each experiment were pooled during protein extraction, so that we analysed a total of 24 cellular proteomes.

**Figure 1.**
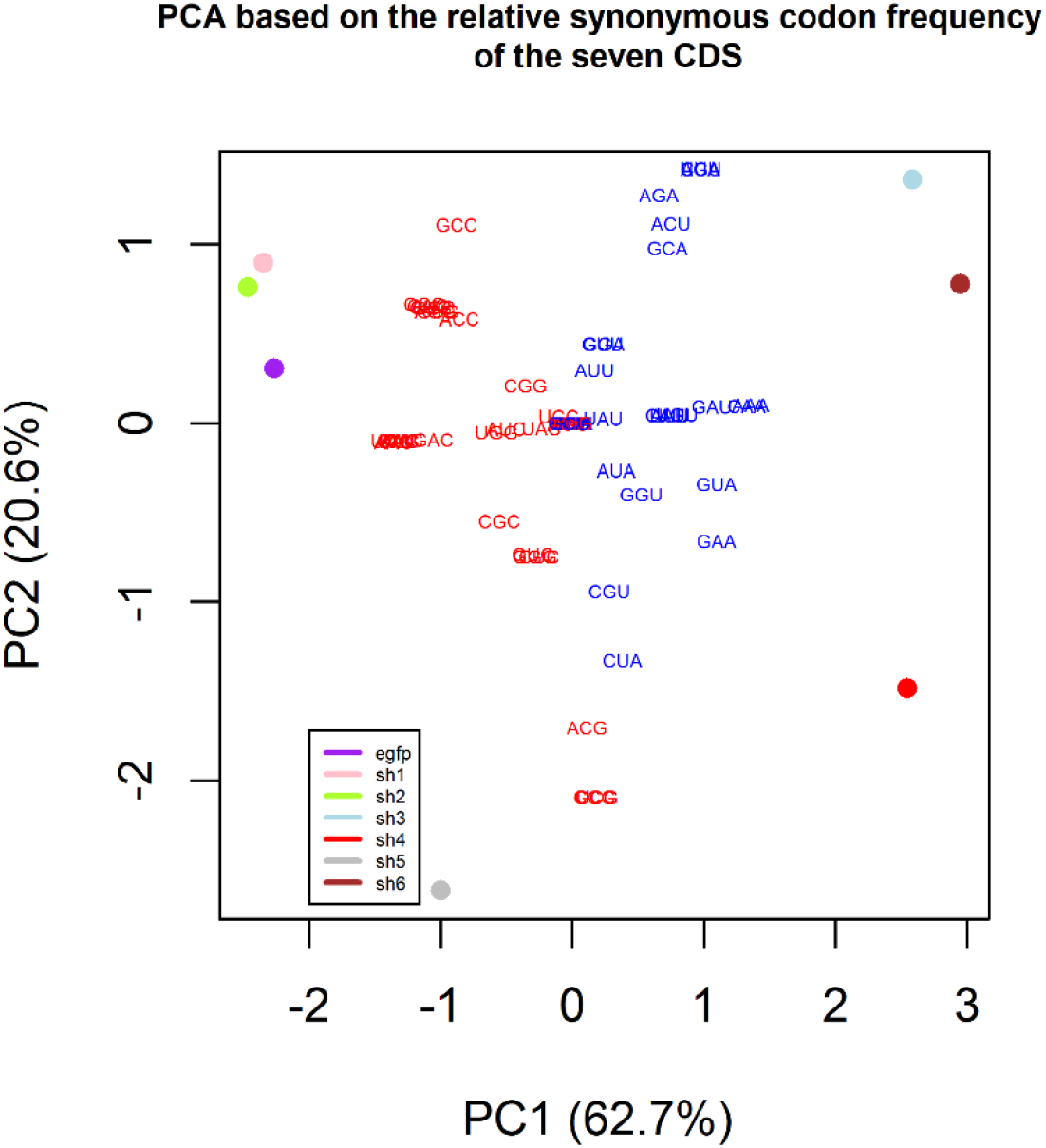
Panel A. Principal Component Analysis based on the relative synonymous codon frequency (RSCF) of the different plasmidic sequences designed. For each seven CDS (the six synonymous ones of the *shble* gene and the one of the *egfp* gene), the profile of their 59 RSCF values (see Methods) were used to perform a PCA. Over-humanized sequences (*i.e*., those rich in human most favored codons: Shble#1 in pink, Shble#2 in green and egfp in purple) cluster together on the top-left of the PC1/PC2 space, while sequences enriched in human rare codons occupy other extreme parts of the PC1/PC2 space (Shble#3 in light blue, Shble#4 in red, Shble#5 in grey, Shble#6 in brown). RSCF profiles allows sequence discrimination based on CUBias. Loadings of each of the 59 amino acids-encoding codons are plotted and displayed with the following color code: those ending either with U or A are indicated in blue and those ending either with C or G are indicated in red.

A principal component analysis performed on our transcriptomic data (exclusively using cellular mRNAs, *i.e*., heterologous mRNAs excluded) revealed that the main source of variation across our samples stems from transfection itself (Fig. S1). The first principal component captured 67% of the total transcriptomic variance, with Mock samples displaying values divergent from all other versions. Importantly, variation along the first component was largely explained by variation in the total amount of heterologous transcripts in the sample (**Fig. 1B**, Spearman correlation coefficient: 0.87, P = 4.8e-7). Since heterologous expression was the largest determinant of transcriptomic changes, we focused on the identification of differentially expressed (DiffExp) cellular transcripts compared to the Mock control for each transfected plasmid version. The total number of transcripts identified as DiffExp in at least one condition was 1,425 (Table S1). Transfection with our Empty control resulted in 645 DiffExp transcripts, while the number of DiffExp transcripts detected in our experimental conditions varied between 487 for Shble#6 and 1,311 for Shble#3, as a function of heterologous gene expression (Fig. S2). Most of the 645 DiffExp mRNAs in the Empty control were also identified as DiffExp in all six (64%) or in five out of the six (82%) *shble* synonymous versions (**Fig. 1C**). The precise overlap between DiffExp mRNAs in the Empty and in each of the six *shble* versions is given in Fig. S3. Finally, and independently of their behaviour in the Empty condition, we observe that a vast majority of transcripts DiffExp compared to the Mock in each version are shared among several *shble* synonymous versions, despite the very different CUBias of the recoded gene (Fig. S4). We hence interpret that our sets of DiffExp transcripts constitute a fundamental part of the cellular response to transfection and to heterologous gene expression, and not to a specific *shble* synonymous version. Functional enrichment analysis supported this conclusion, with all seven sets of DiffExp transcripts relative to the Mock sharing the same top three enriched categories, namely *Inflammatory response, Type I interferon signaling pathway* and *Response to virus*. Two other immunity-related categories appeared over-represented in most of the sets of DiffExp transcripts: *Negative regulation of viral genome replication* and *Immune response* categories (Table S2). Of note, an overwhelming majority of mRNAs belonging to these five categories appeared to be up-regulated in comparison to the Mock condition, suggesting transfection triggered an inflammatory response in our HEK293 cells. Overall, the global trends for our transcriptomic results are that: i) transfected cells undergo substantial changes in the transcriptome; ii) the extent of these changes correlates with the extent of heterologous gene expression; iii) the bulk of the transcriptomic response are shared among conditions, irrespective of the CUBias of the transfected *shble* version; and iv) transcriptomic changes upon transfection largely overlap the activation of the cellular response to viral infection and inflammatory response.

**Figure 1.**
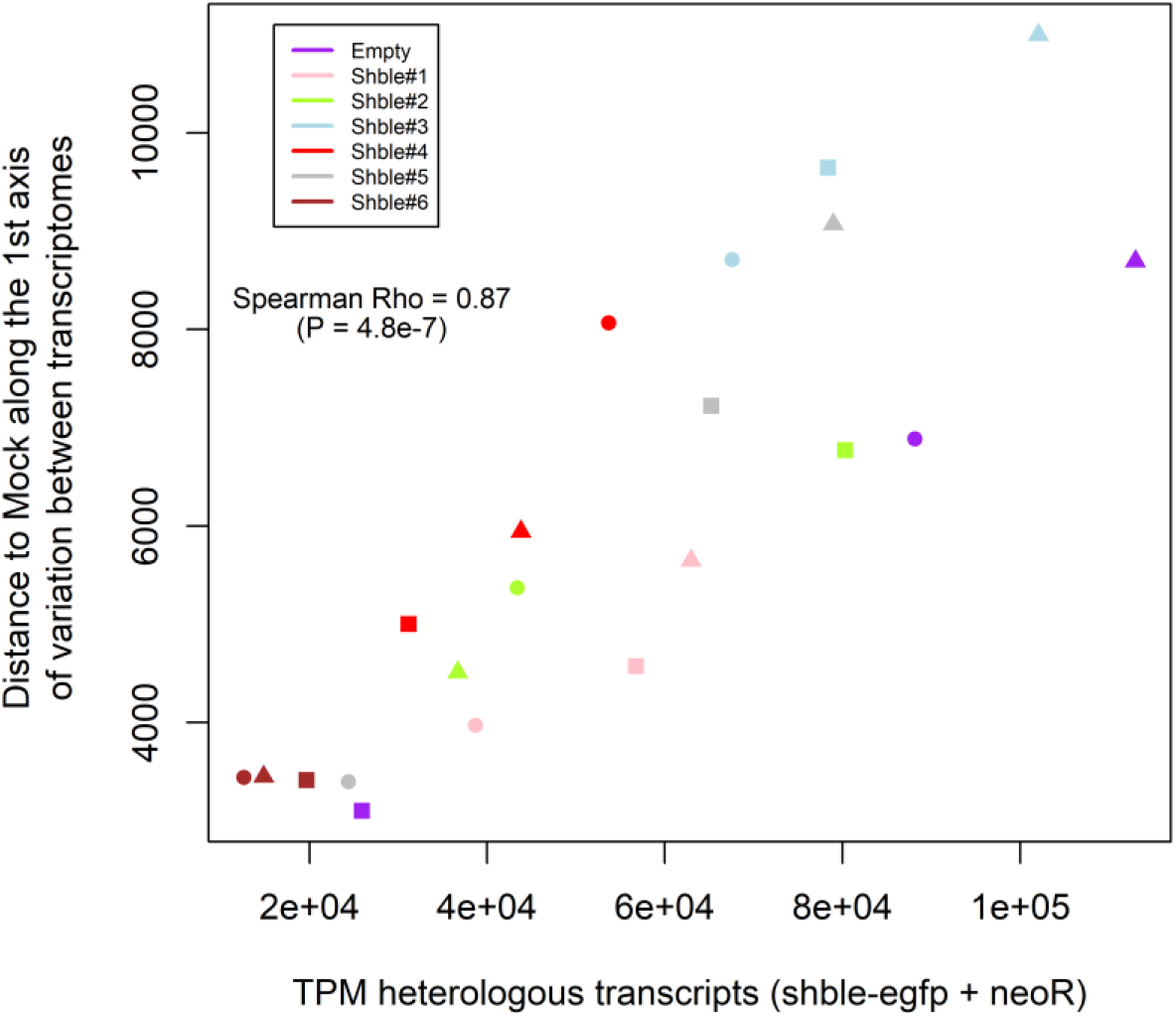
Panel B. Link between the global transcriptome profile relatively to the Mock condition and the quantity of heterologous transcripts expressed. The y-axis represents the distance between transfected and Mock samples, obtained as the difference between their projections onto the first axis of the PCA based on expression of cellular transcripts (Fig. S1). Samples are color-coded according to the different conditions: Empty in purple, Shble#1 in pink, Shble#2 in green, Shble#3 in light blue, Shble#4 in red, Shble#5 in grey, Shble#6 in brown. The three experimental batches are symbolized by different shapes.

**Figure 1.**
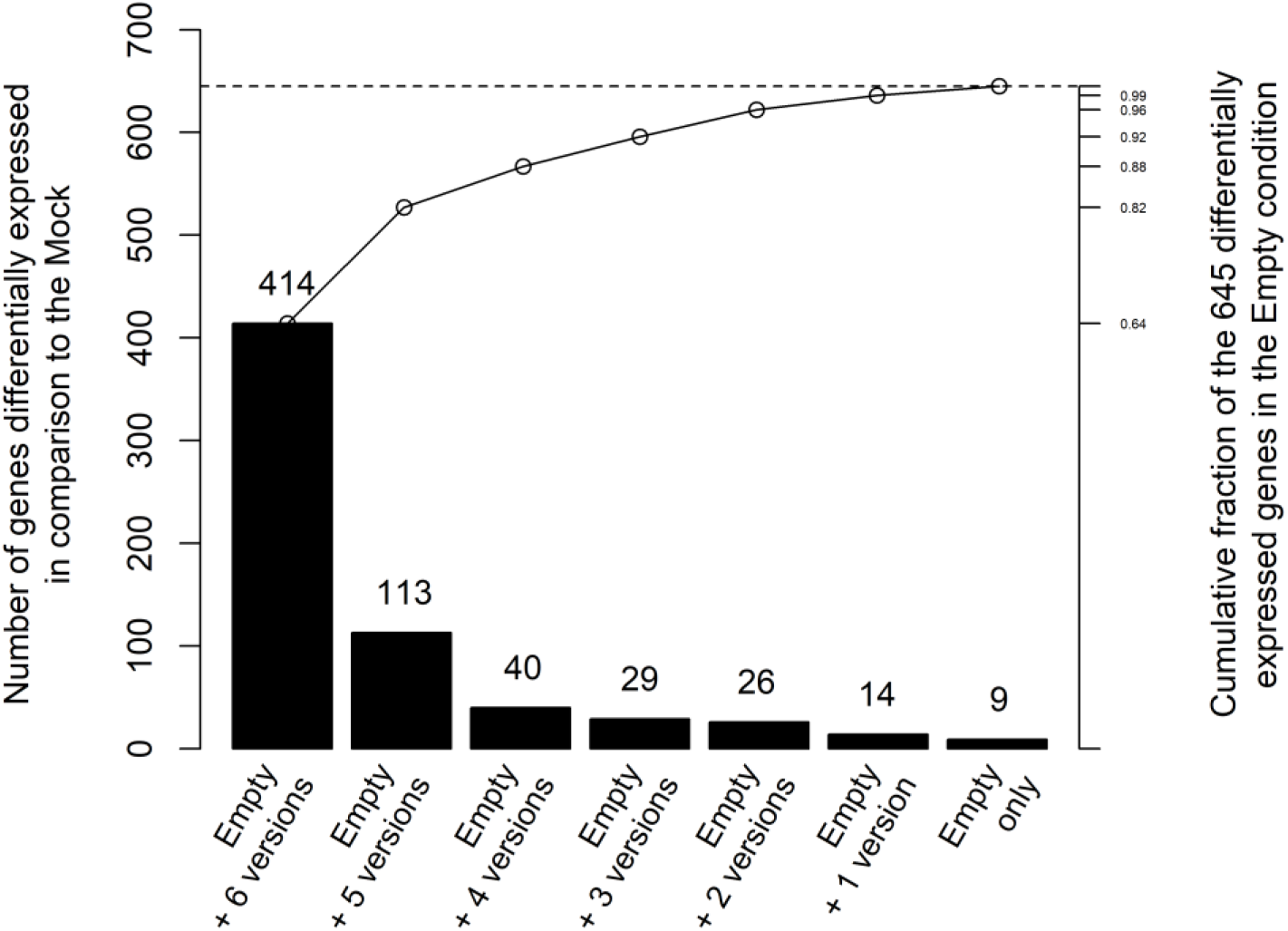
Panel C. Match between differentially expressed (DiffExp) transcripts in the Empty control (only *shble* expressed) and DiffExp transcripts in the six *shble-egfp* synonymous versions. Each of the 645 DiffExp transcripts in the Empty condition were assigned to a category depending on the number of other conditions this gene was found to be DiffExp. In all cases the transcripts are labelled as DiffExp with respect to the Mock control condition. Bar heights correspond to the number of genes identified in each category (left axis). The secondary axis on the right reports the cumulative fraction of the 645 DiffExp transcripts in the Empty control condition as each category is added to the previous one. Data should be read as follows, using “Emtpy + 5 versions” as an example: among the 645 transcripts DiffExp between the Empty version and the mock, a total of 414+113 transcripts (*i.e*., 82% of the 645 transcripts – right axis) are also DiffExp in five or more of the *shble-egfp* synonymous conditions.

### Lack of directional changes in the cellular proteome upon transfection and heterologous gene expression, irrespective of the codon usage bias of the heterologous genes

Our experimental setup of paired transcriptomic and proteomic data allowed us to estimate intra-sample covariation between mRNA and protein levels. Overall, we detected a total of 4,005 protein entries, with an average of 2,723 proteins detected per sample, and 2,891 proteins detected in all three replicates of at least one condition (Table S3). Distribution of relative intensity-Based Absolute Quantification (riBAQ) values for proteins detected in each sample is given in Fig. S5. When coupling proteomic data with transcriptomic data, we observed that variation in mRNA levels accounts for around one-third of the variation in protein levels (median R^2^=0.31, min=0.29, max=0.35, Table S4). A representative example is provided in Fig. S6. These values agree with previous measurements performed on mammalian cells, with studies reporting correlation values around 0.40 for mouse cells (**Schawanhaüsser et al, 2011**) and between 0.30 and 0.40 for human cells (reviewed in **Vogel & Marcotte, 2012**).

We next used proteomic data to investigate whether we could find evidence for a shared response across versions, similar to the one described above at the transcriptomic level, or alternatively whether a *shble* synonymous version-specific response is observed after transfection at the protein level. The two first axes of a PCA performed on our proteomic data (exclusively using cellular proteins) captured an important fraction (77%) of the global proteome variation among samples (Fig. S7). However, in contrast with variation at the transcriptomic level, the experimental conditions did not spread following any evident pattern. Especially, variation along the first component was not driven by variation in the total amount of heterologous transcripts expressed by the samples (Spearman correlation coefficient = 0.099; P = 0.67). The absence of clustering in the PCA was still notable when considering the set of 2,891 proteins that were robustly detected in all the three replicates of at least one condition (data not shown). Accordingly, differential expression analysis failed to identify differentially expressed proteins, either with regards to transfection itself or regarding the different *shble* synonymous versions in the transfected plasmids. We tried to refine this analysis by defining a set of proteins that could maximize the chances of detecting DiffExp proteins between conditions. We identified thus all proteins that were detected in all three replicates of a same condition and that were not detected in any of the three replicates of at least one other condition (see Methods and Table S3), reasoning that if an effect was to be found, working specifically on this subset of proteins would maximize variation across versions. We found 369 proteins that fulfilled this stringent criterion, but again a Principal Component Analysis failed at identifying systematic differences between conditions (Fig. S8). Overall, we concluded that in our experimental setup, the cellular proteomic response to heterologous expression was much less important than the transcriptomic one.

### High translation level of heterologous sequences does not impair translation of cellular genes with a similar codon usage bias

For prokaryotic and unicellular eukaryotic systems it has been proposed that overexpressing heterologous genes with extreme CUBias can affect the translation efficiency of other genes (**Kudla et al, 2009; Shah et al, 2013; Frumkin et al, 2018**). To test whether this was also the case in our human cells, we chose as a proxy for translation efficiency of a given gene the ratio protein-over-mRNA (expressed as an riBAQ/TPM ratio). We first studied the impact of heterologous expression on the translation efficiency of cellular genes by comparing the Empty condition to the Mock condition. Cells under the Empty condition were transfected with a plasmid encoding only the *egfp* gene, whose sequence, “enhanced” for expression in human cells, is strongly biased towards the use of the most frequent codons in the human genome (COUSIN value of *egfp:* 3.38). This first level of analysis hence provide insights into potential consequences of over-expressing an “over-humanized” gene on the translation efficiency of cellular genes. Changes in the riBAQ/TPM ratio in Empty *vs*. Mock samples were calculated for 2,471 cellular genes and are depicted as a function of the CUBias of the corresponding gene (**Fig. 2A**). Our results show that the intensity of changes in riBAQ/TPM for cellular genes when EGFP is actively expressed is not a function of gene’s CUBias (slope = 0.00378, F-test P = 0.29). We performed the same analysis comparing riBAQ/TPM ratios in Shble#1 and Shble#2 conditions against the Mock. These two versions encode *shble* using an over-humanized CUBias, in the same direction than *egfp* (COUSIN values to the human genome for Shble#1 and Shble#2 are of 3.47 and 3.42, respectively). Again, we do not observe any trend in protein-over-mRNA levels as a function of the CUBias of the corresponding gene (**Fig. 2B,** slope = 0.0048, F-test P = 0.18 and **Fig. 2C**, slope = −0.0059, F-test P = 0.11). These results go in the same direction as the ones presented in Fig. 2A and do not support the hypothesis that translation of highly expressed over-humanized genes especially impairs translation of cellular genes also enriched in these human-frequent codons.

**Figure 2.**
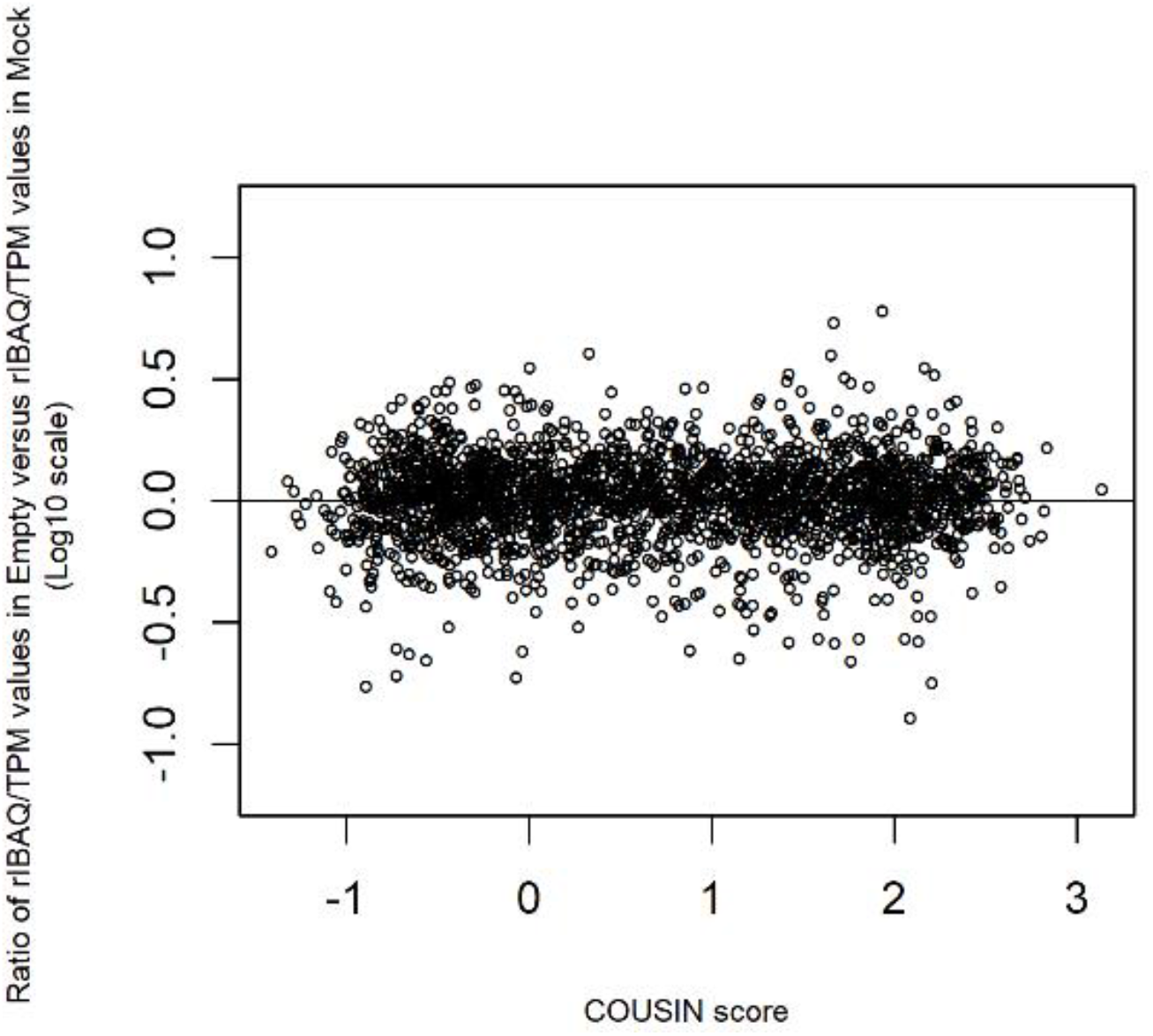
Panel A. Protein-over-mRNA ratios in the Empty condition compared to the Mock condition as a function of individual gene’s codon usage bias (CUBias). Transcript levels were estimated as Transcripts Per Million (TPM), while protein levels were estimated as relative intensity-Based Absolute Quantification (riBAQ). Each dot represents, for one gene, the averaged [riBAQ/TPM]_Empty_ normalized by the [riBAQ/TPM]_Mock_, calculated from the n=3 transfection experiments. This ratio of ratios reflects the extent to which cellular transfection and heterologous expression of a gene with an over-humanized sequence (*egfp*, encoded in the Empty condition) affects translation efficiency of cellular transcripts. This ratio has been calculated for the 2,471 genes for which both proteomic and transcriptomic values were available. For visual purposes, the horizontal line centered on the y=0 value (log scale) is shown. The x-axis displays the match between the CUBias of each individual gene to that of the human genome average, calculated using the COUSIN index: negative scores correspond to genes with CUBias opposite to the human average while scores above one correspond to genes with CUBias similar in direction but of stronger intensity than the human average. The COUSIN value of the *egfp* expressed in the Empty condition is of 3.38. No link was found between gene’s CUBias and the gene’s [riBAQ/TPM]_Empty_ ratio normalized by the [riBAQ/TPM]_Mock_ ratio.

**Figure 2.**
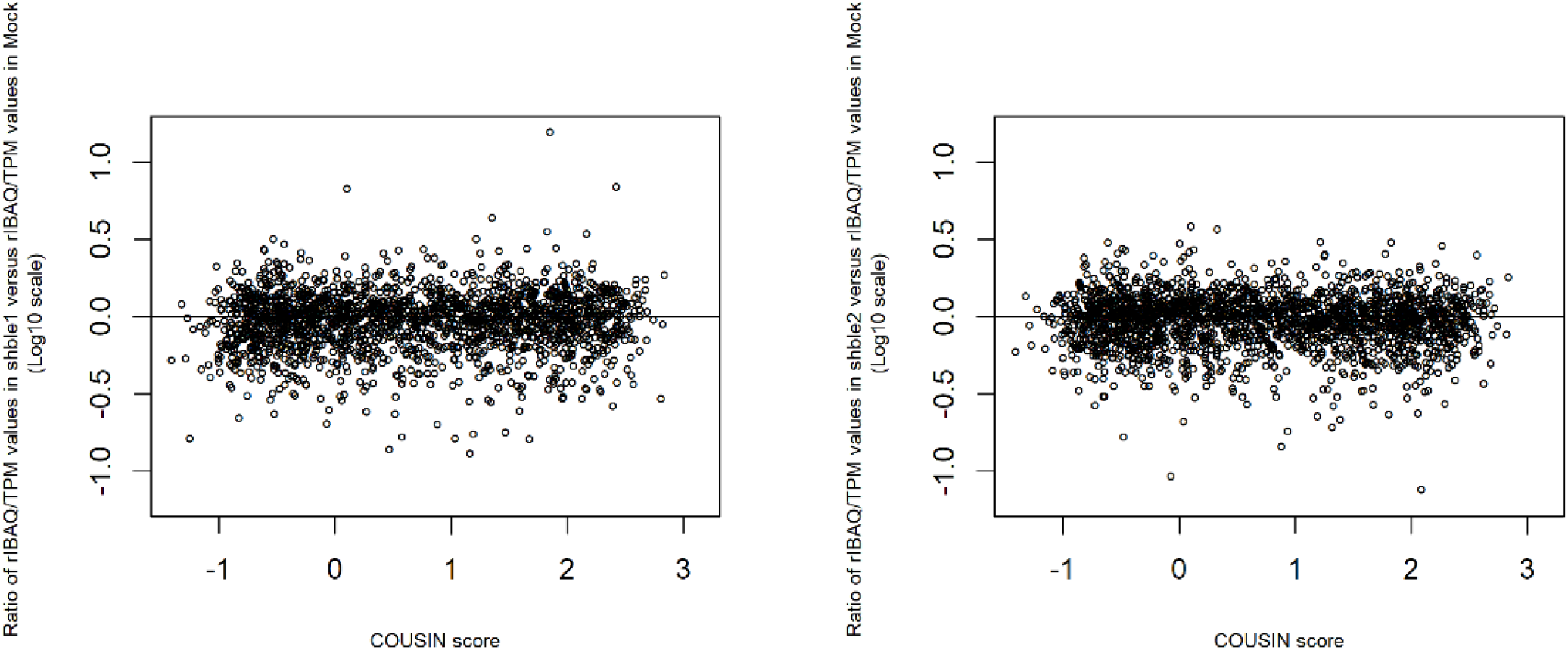
Panel B and C. Similar to Figure 2.A, but for the Shble#1 condition compared to the Mock (Panel B, left) or for the Shble#2 condition compared to the Mock (Panel C, right). The *shble* CDS in the Shble#1 version had been synonymously recoded using systematically for each amino acid the most used among the synonymous codons in the human genome average. It corresponds thus to an over-humanized heterologous gene (COUSIN value of this version =3.47). The *shble* CDS in the Shble#2 version had been synonymously recoded using systematically for each amino acid the GC-richest among the two most used synonymous codons in the human genome average (COUSIN value of this version = 3.42). It differs only in eight codons from the Shble#1 version. As in Figure 2.A, no link was found between gene’s CUBias and the gene’s riBAQ/TPM ratio of ratios, neither for Shble1 versus Mock (left), nor for Shble#2 versus Mock (right).

To extend the analysis of the impact of heterologous gene expression on translation of cellular genes to the range of CUBias explored in our setup, we aimed at identifying cellular genes that presented consistent trends of changes in protein-over-mRNA ratio *across* conditions. As a proxy for translation level of the heterologous genes we used the total amount of EGFP and SHBLE detected in each sample, using the sum of the corresponding riBAQ values. We stratified the experimental conditions in two mutually exclusive sets: those that expressed heterologous genes enriched in codons frequently used in the human genome (*i.e*., Empty, Shble#1 and Shble#2) and those that expressed heterologous genes enriched in codons rarely used in the human genome (*i.e*., Shble#3, Shble#4, Shble#5 and Shble#6).

We first focused on conditions using human-frequent codons (Empty, Shble#1 and Shble#2), exploring correlation between variation in riBAQ/TPM ratios for each individual gene and variation in the total amount of heterologous proteins. This analysis used twelve underlying data points per gene as explanatory variable: three values for the MOCK control samples and nine values for the transfected samples. Out of the 2,550 cellular genes that we could analyze, 235 displayed a significant co-variation between riBAQ/TPM ratios and heterologous protein levels: 109 genes showed a positive and 126 showed a negative, significant co-variation (**Fig. 3A**, yellow and green sets, respectively; Table S5). Results for a representative example of a gene with a negative association (green set) are given in **Fig. 3B** (left, P = 0.0092). We next compared these two sets of genes in terms of CUBias, with the hypothesis that a competition for translational resources between cellular and heterologous transcripts should primarily negatively affect translation of genes enriched in most human-frequent codons (*i.e*., in the same codons used by *egfp* and Shble#1 and Shble#2 versions). Instead, we found that the median COUSIN score for the two gene sets were not significantly different (0.58 for the set of 126 genes; 0.45 for the set of 109 genes; Wilcoxon Mann-Whitney test P = 0.58, **Fig. 3C**). An additional Anderson-Darling test failed to reject the null hypothesis that the COUSIN value distributions for each dataset could have been drawn from a same underlying distribution and were thus not significantly different (P=0.33, **Fig. 3C**). Finally, these two gene sets, with translation positively or negatively affected by overexpression of heterologous genes enriched in human-frequent codons, do not display different distribution of COUSIN values than the overall cellular genes (Fig. S9). These results confirm the trend communicated above (Figure 2) and show that cellular genes whose translation was positively or negatively impacted by overexpression of heterologous genes enriched in human-frequent codons do not display different CUBias: high expression of heterologous genes enriched in human-frequent codons does not impose a substantial differential burden on host translation efficiency of cellular genes as a function of their CUBias.

**Figure 3.**
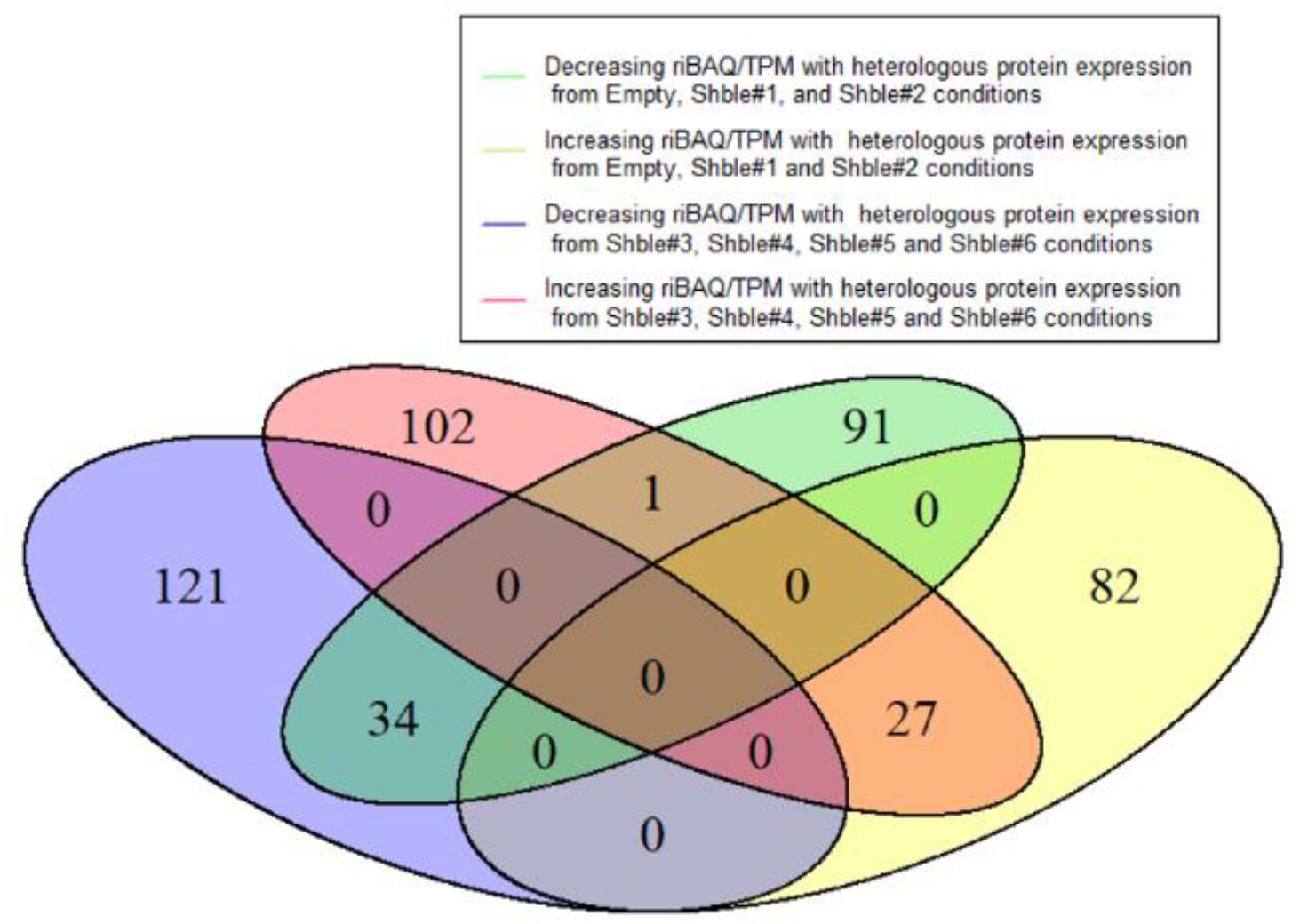
Panel A. Categories of genes according to their variation in riBAQ/TPM ratios with varying heterologous protein expression across conditions. The ratio riBAQ/TPM was taken as a proxy for the translation efficiency of a given transcript. The green set corresponds to cellular genes displaying a decreasing translation efficiency as heterologous protein expression under the conditions Empty, Shble#1 and Shble#2 increases (*i.e*., when expressing humanized heterologous genes increases), while the yellow set corresponds to genes displaying an increasing translation efficiency under these same conditions. The blue set corresponds to cellular genes displaying a decreasing translation efficiency as heterologous protein expression under the conditions Shble#3, Shble#4, Shble#5 and Shble#6 increases (*i.e*., when expressing under-humanized heterologous genes increases), while the pink set corresponds to genes displaying an increasing translation efficiency under these same conditions. Note that, by construction, neither the green and yellow sets nor the blue and pink sets overlap.

**Figure 3.**
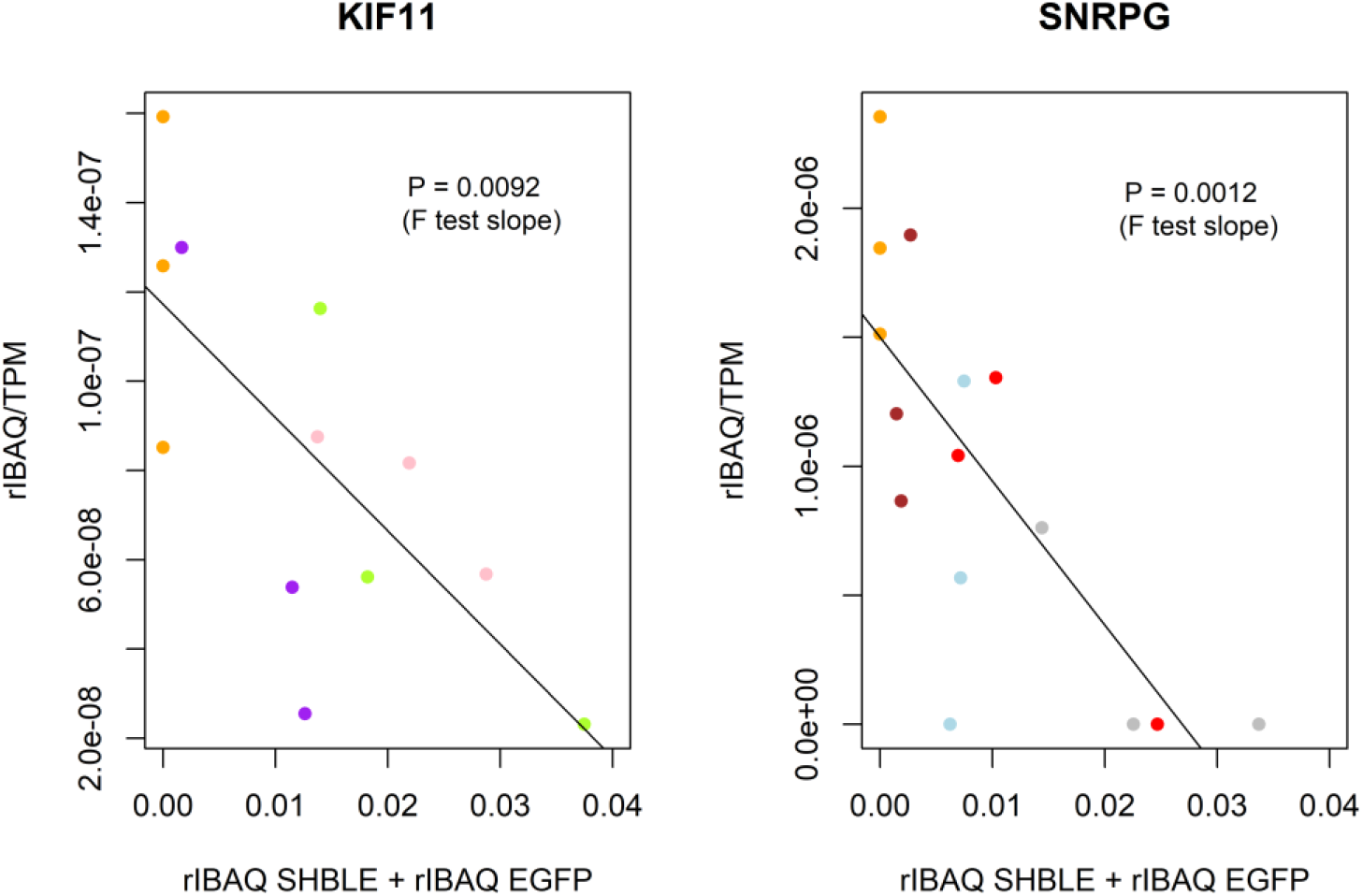
Panel B. Example of two cellular genes displaying significant variation in their riBAQ/TPM ratio as a function of heterologous protein expression levels across samples. The ratio riBAQ/TPM was taken as a proxy for the translation efficiency of a given transcript. KIF11 (left) belongs to the green set shown in Figure 3.A (negative association with increasing amount of protein expressed from over-humanized heterologous genes); SNRPG (right) belongs to the blue set shown in Figure 3.A (negative association with increasing amount of protein expressed from non-humanized heterologous genes). The x-axis represents heterologous proteins expression in the different samples (n=3 by condition). Samples are color-coded according to the transfected construct: Shble#1 in pink, Shble#2 in green, Shble#3 in light blue, Shble#4 in red, Shble#5 in grey, Shble#6 in brown, Empty in purple and Mock in orange. P-values of F-tests testing the significance of regression slopes are indicated.

**Figure 3.**
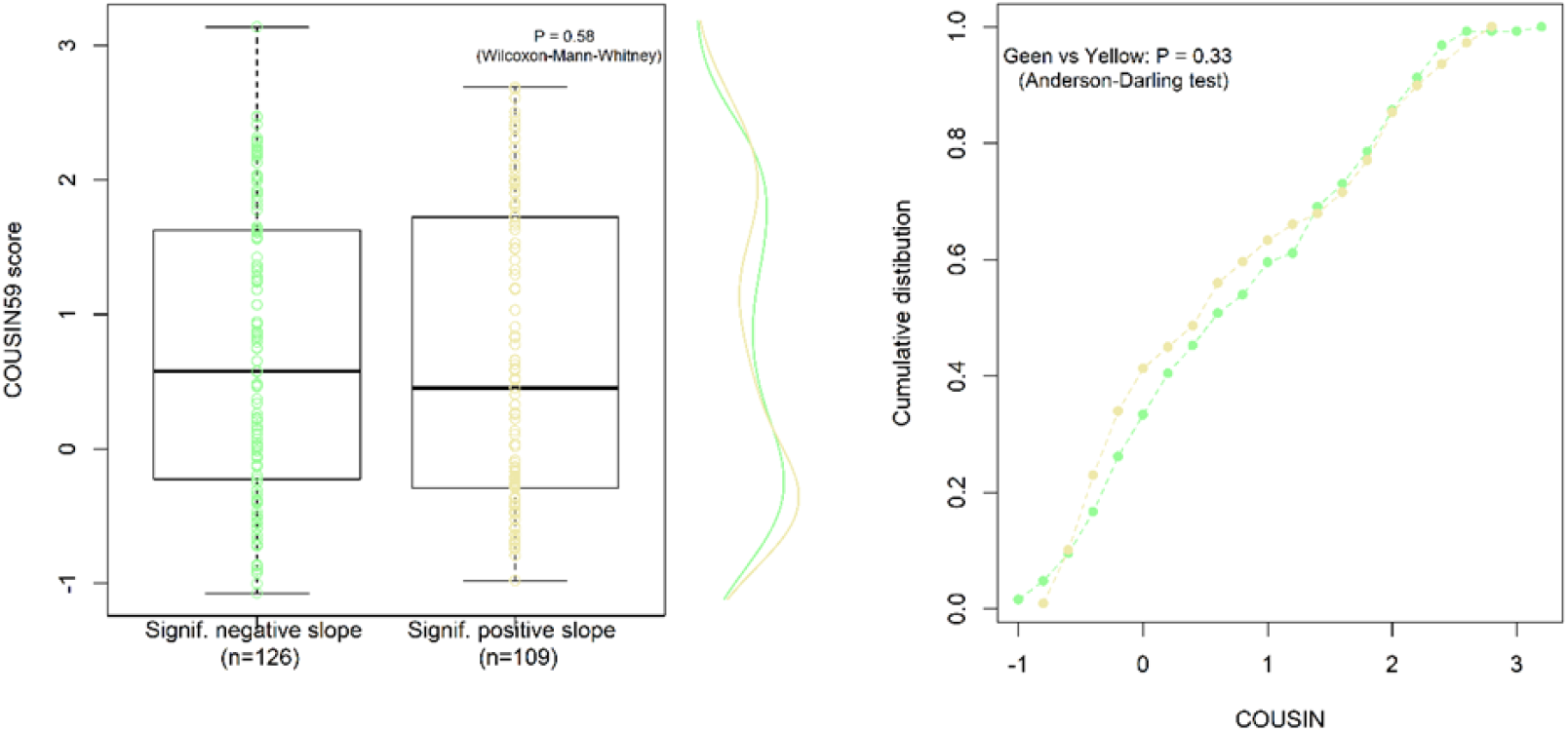
Panel C. Left: COUSIN scores for cellular genes displaying either negative (green) or positive (yellow) changes in the riBAQ/TPM ratio values under increasing expression of over-humanized genes (Empty, Shble#1 and Shble#2). Below each boxplot the underlying number of genes included is shown. The P-value of the Wilcoxon-Mann-Whitney test contrasting median COUSIN values is indicated. Marginal distributions of COUSIN values underlying each boxplot are shown on the right y-axis. Right: Cumulative distribution of COUSIN values for the two sets of genes described in the left. P-values for of the Anderson Darling test contrasting the two distributions is indicated.

It is conceivable that we failed to identify a differential impact of over-humanized protein expression on host translation efficiency due to a pool of tRNAs for the considered codons large enough to accommodate the demand exerted by translation of heterologous mRNAs. Alternatively, we can expect the pool of tRNAs decoding human rare codons to be more limiting and therefore more prone to be depleted by the demand imposed by the translation of heterologous mRNAs rich in such rare codons. We thus perform our analysis with the second set of conditions, focusing on conditions expressing heterologous genes enriched in codons underrepresented in the human genome. This analysis used 15 underlying data points per gene as explanatory variable: three values for the Mock control samples and twelve values for the transfected samples (*i.e*. Shble#3, Shble#4, Shble#5 and Shble#6). Out of the 2,580 cellular genes that we could analyze, 285 displayed a significant co-variation between riBAQ/TPM ratios and heterologous protein levels: 130 genes showed a positive and 155 showed a negative, significant co-variation (**Fig. 3A**, pink and blue sets, respectively; Table S5). Results for a representative example of a gene with a negative association (blue set) are given in **Fig. 3B** (right, P = 0.0012). We compared then the CUBias of the genes in each of these two sets and we observed that they differ in their match to the average human CUBias: cellular genes showing a negative association with overexpression of heterologous genes enriched in human-rare codons displayed higher COUSIN scores than genes displaying positive association (respective median values 0.79 and 0.13; Wilcoxon Mann-Whitney test P = 0.010, **Fig. 3D**). The distribution of COUSIN values between the two gene datasets is also significantly different (Anderson-Darling test, P = 0.0010, **Fig. 3D**). Furthermore, cellular genes negatively affected by overexpression of heterologous genes enriched in human-rare codons do not display different COUSIN values distribution than the ensemble of the cellular genes, while genes positively affected do display a different CUBias distribution that is shifted towards lower COUSIN values (Fig. S10). These results are counter-intuitive because – assuming cellular mRNAs compete with heterologous mRNAs for access to rare-codon decoding tRNAs – we would instead have expected cellular genes negatively impacted to be the ones preferentially using these non-optimal codons that are required for translating heterologous transcripts and hence to be those presenting lower COUSIN values.

**Figure 3.**
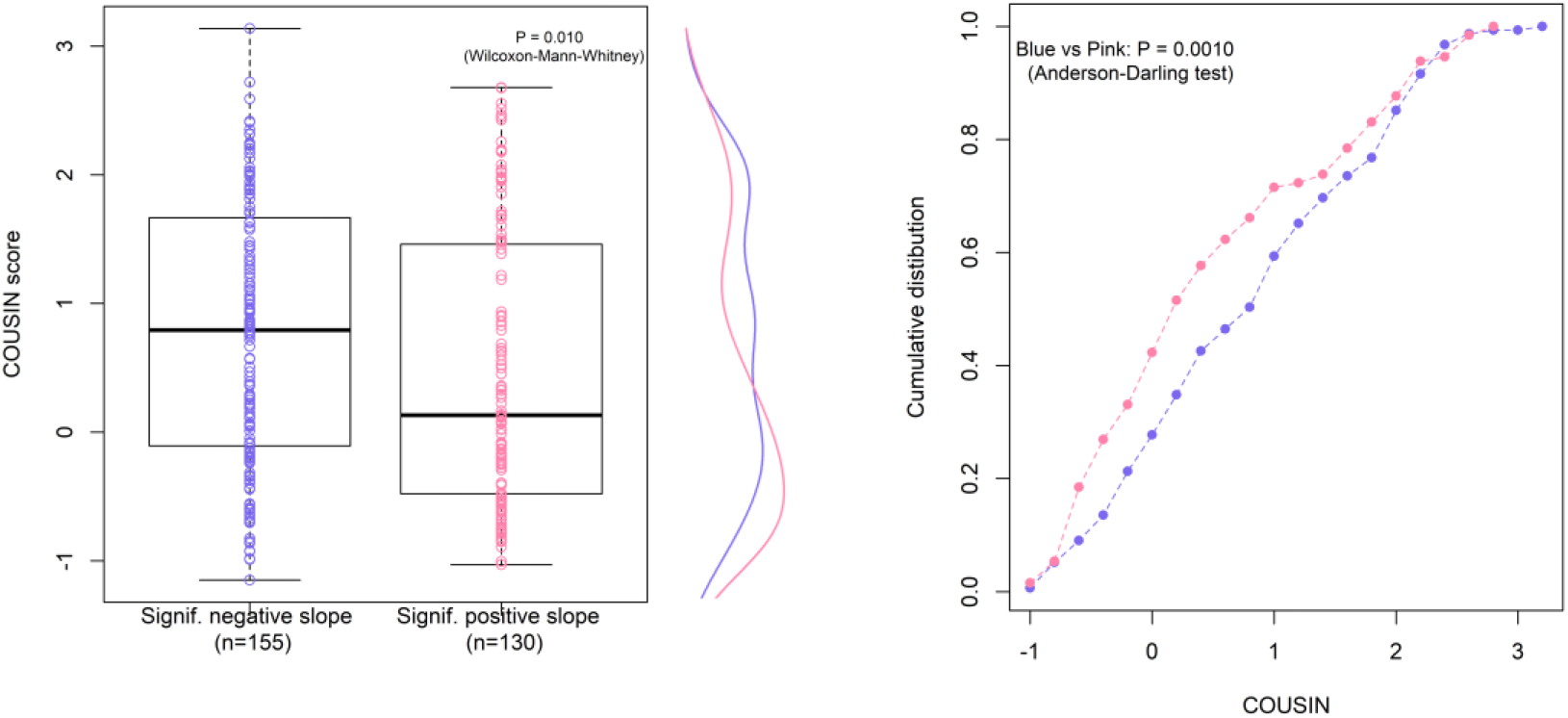
Panel D. Left: COUSIN scores for cellular genes displaying either negative (blue) or positive (pink) changes in the riBAQ/TPM ratio values under increasing expression of non-humanized genes (Shble#3, Shble#4, Shble#5 and Shble#6). Below each boxplot the underlying number of genes included is shown. The P-value of the Wilcoxon-Mann-Whitney test contrasting median COUSIN values is indicated. Marginal distributions of COUSIN values underlying each boxplot are shown on the right y-axis Right: Cumulative distribution of COUSIN values for the two sets of genes described in the left. P-values of the Anderson Darling test contrasting the two distributions is indicated.

For the sake of completeness, we repeated all these analyses (*i.e*., using the two mutually exclusive sets of conditions), after having excluded cellular genes that were significantly affected in the same direction in both datasets (*i.e*., after having removed the 27 cellular genes that are positively affected in both cases and the 34 cellular genes that are negatively affected in both cases). The results did not change with regards to those communicated above (Fig. S11).

Overall, the global trends for our experiment results does not support thus the hypothesis for resource competition among mRNAs for the rare cellular resources of the translation machinery, as overexpression of genes with a given CUBias does not hinder translation of other genes with similar CUBias.

### Heterologous expression of genes with extreme CUBias does not lead to a global alteration of the translation efficiency of cellular transcripts

We finally used a more integrative approach to examine how the global codon repertoire present at the transcriptomic layer ‘flows’ to the proteomic layer. This approach follows a supply-and-demand reasoning: codons present in the pool of cellular mRNAs exert a demand for being decoded whereas the codon composition inferred from the cell proteome informs us to what extent this demand has been met. For this purpose, we calculated for every sample the occurrence of each codon by accounting for their relative abundance, as they are represented in the whole cellular transcriptome, in the whole cellular proteome, or in the set of transcripts that matched detected proteins (Table S6). We next computed the relative synonymous codon frequency (RSCF) from these ponderated codon counts (see Methods) and recapitulated the obtained RSCF profiles in the PCA shown in **Fig. 4A**. To avoid introducing bias due to difference in sensitivity of features detection (~ 14,000 mRNAs detected versus ~ 3,000 proteins detected) when comparing proteome RSCF and transcriptome RSCF, we decided to use exclusively the set of transcripts that matched detected proteins instead of the whole transcriptome data for the rest of the analysis. By dividing proteome RSCF of each codon by its transcriptome RSCF counterpart, we obtained for each sample an integrated view of how content of each codon changed between mRNAs present in the cell and mRNAs that had been translated into proteins. We called this variable Prot-to-RNA RSCF and considered it as a proxy of whole-cell translation efficiency of codons. The resulting variation across samples for their Prot-to-RNA RSCF vectors is shown in **Fig. 4B**. We observed that Shble#3 seem to have preferentially translated GC-ending codons. More generally, we observed that most AT-rich versions (Shble#3, Shble#4 and Shble#6) have positive scores on PC2, while most GC-rich versions (Shble#1, Shble#2, Shble#5 + Empty) have negative scores. We checked the correlation between heterologous transcripts expression and both PC1 (62%) and PC2 (15%) of the variation of Prot-to-RNA RSCF across samples, to see if it drives some of the variation and hence could cofound the effect of our synonymous recoding of *shble* versions *per se*. None of the correlation was significant, but for PC2 there was a positive trend (Fig. S12). For this reason, we examined the relationship between *shble* sequence features and PC2 of Prot-to-RNA RSCF, after having removed the effect of heterologous expression (see Methods). We focused on *shble* CDS GC-richness and COUSIN score and found a significant and negative correlation for GC-richness (R^2^ = 0.28, P = 0.015 - **Fig 4C**). Since higher PC2 scores represent samples that favored the translation of GC-ending codons (Fig. 4B), the result of **Fig. 4C** suggests a differential effect in terms of codon third base-dependent global translation efficiency in host cells, driven by base composition of the synonymous version they express. Samples expressing GC-depleted versions of *shble* (such as Shble#3) preferentially translate GC-ending codons, whereas samples expressing GC-rich versions of *shble* (such as Shble#1 or Shble#2) preferentially translate AT-rich codons. However, this effect is subtle since it only emerges when looking at PC2, which captures only 15% of the total variation across samples in their whole-cell translation efficiency.

**Figure 4.**
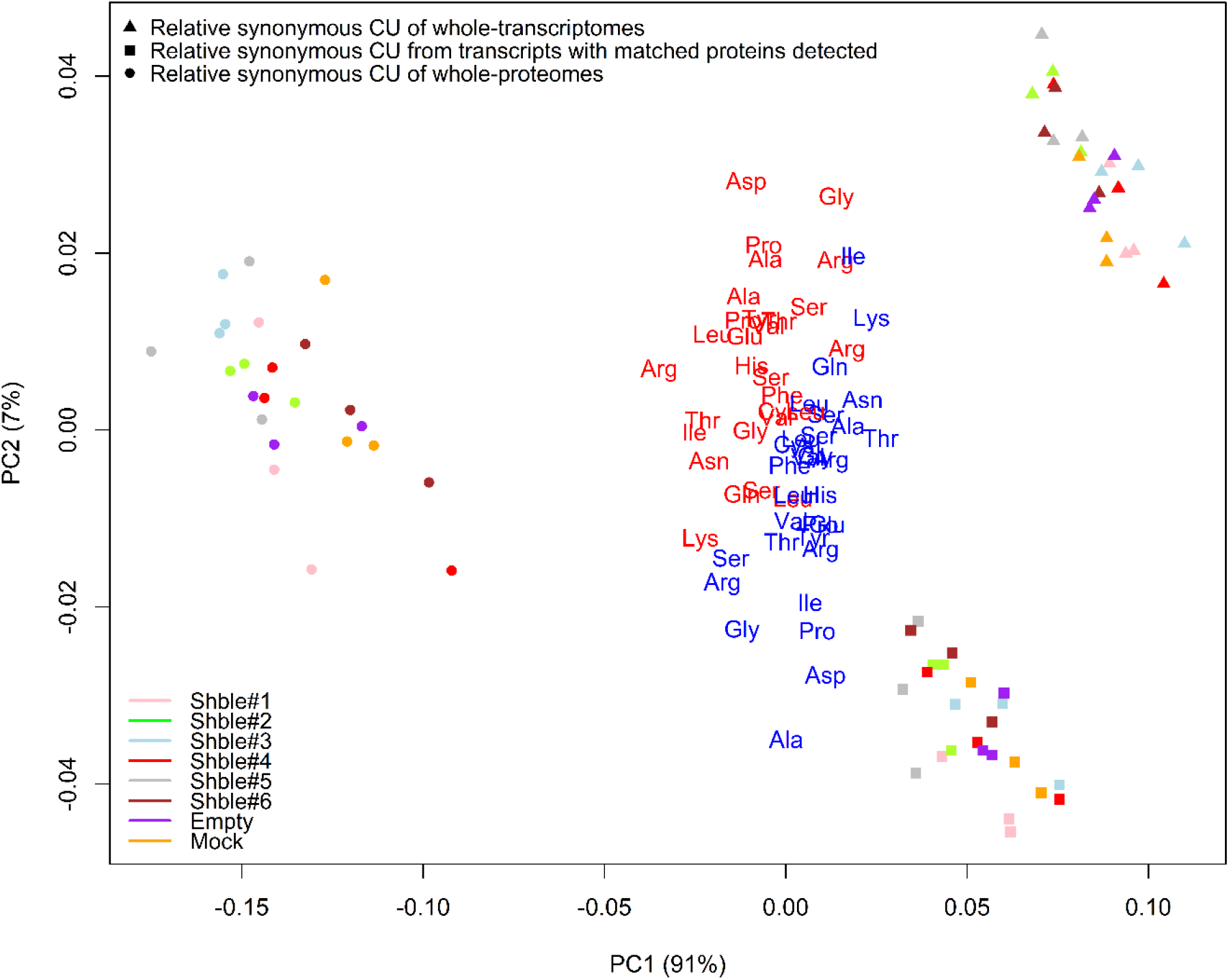
Panel A. Relative synonymous codon (RSCF) frequency profiles from different datasets of expression. For each sample, a vector of RSCF has been obtained using codon content ponderated by codons expression from either transcriptomic or proteomic data. Symbol shapes indicates the expression matrix used to compute these vectors: whole-transcriptome data (triangles), whole-proteome data (dots), or expression of transcripts that match detected proteins (squares). Samples are color-coded according to the transfected construct: Shble#1 in pink, Shble#2 in green, Shble#3 in light blue, Shble#4 in red, Shble#5 in grey, Shble#6 in brown, EMPTY in Purple and Mock in orange. Eigenvectors loads of the covariance matrix are superimposed to the graph and are colored according to the nucleotide composition at the codon 3^rd^ position (blue for A or U-ending codons, red for G or C-ending codons). Values in parentheses along axes represent the fraction of the total variance captured by the corresponding axis.

**Figure 4.**
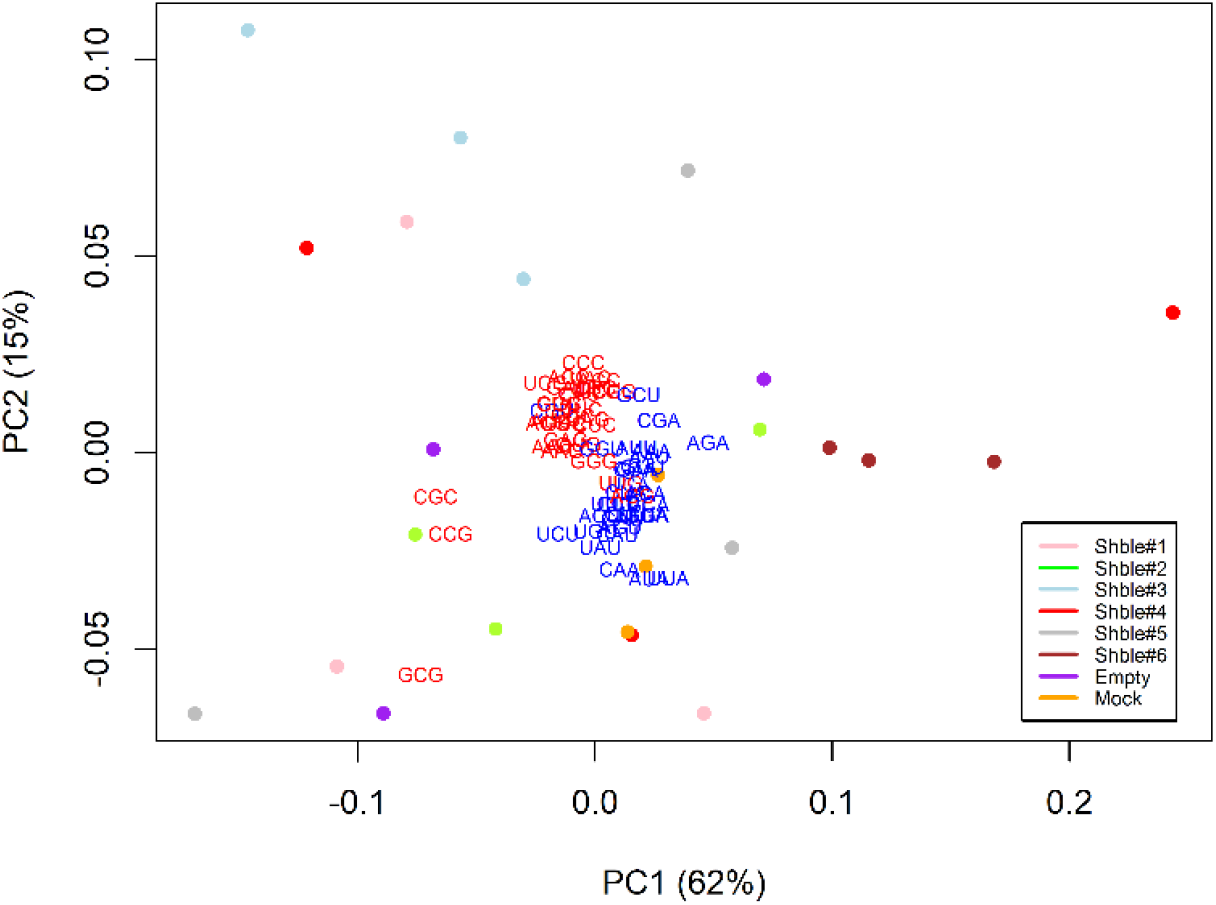
Panel B. Inter-sample variation of Prot-to-RNA RSCF profiles. The Prot-to-RNA RSCF metrics has been computed using exclusively transcripts expression data from transcripts that matched protein detected by LC-MS/MS (*i.e*., we did not consider whole-transcriptomes). The first two axes of variation from a PCA performed on Prot-to-RNA RSCF profiles using 59 amino acids-encoding codons are shown. Samples are color-coded according to the transfected construct: Shble#1 in pink, Shble#2 in green, Shble#3 in light blue, Shble#4 in red, Shble#5 in grey, Shble#6 in brown, Empty in Purple and Mock in orange. Eigenvectors loads of the covariance matrix are superimposed to the graph and are colored according to the nucleotide composition at the codon 3^rd^ position (blue for A or U-ending codons, red for G or C-ending codons). Values in parentheses represent the fraction of the total variance captured by the corresponding axis.

**Figure 4.**
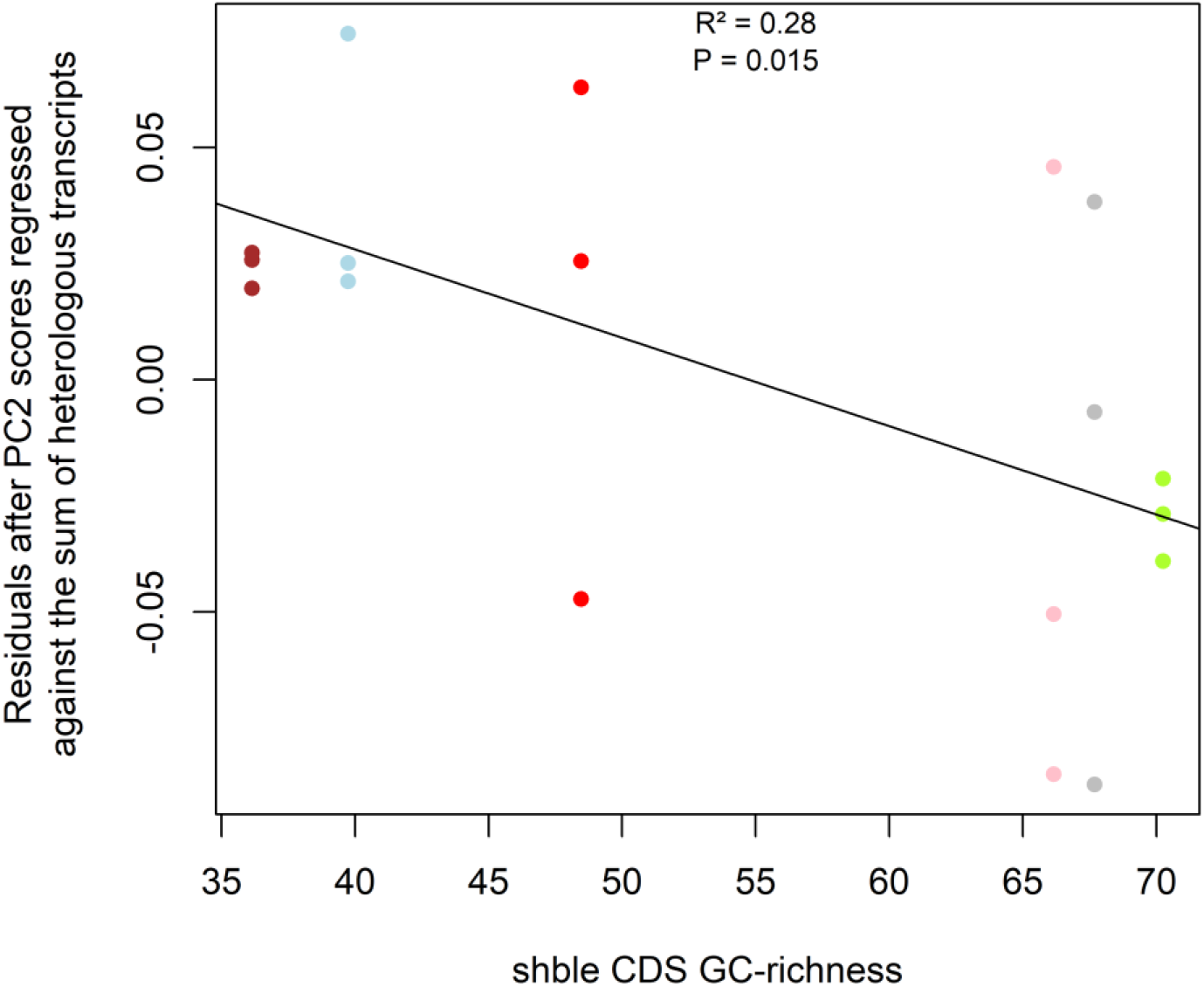
Panel C. Link between base composition of the *shble* version and preferentially translated codons. The y-axis corresponds residuals obtained after samples’ PC2 scores (capturing 15% of the total variance in Prot-to-RNA RSCF, Panel B) were regressed against the sum of heterologous transcripts (Fig. S12). Only the 18 experimental synonymous *shble-egfp* conditions are included. Samples are color-coded according to the transfected construct: Shble#1 in pink, Shble#2 in green, Shble#3 in light blue, Shble#4 in red, Shble#5 in grey and Shble#6 in brown. The x-axis indicates the overall GC-richness of the *shble* CDS of each synonymous version. The regression line is plotted, as well as the P-value of the F-test for the slope significance and the coefficient of determination (R^2^).

## Discussion

We present here a systematic analysis of the changes in the cellular transcriptomic and proteomic profiles upon experimental transfection, using a number of synonymous versions of heterologous genes with divergent CUBias. Our results show that transfection and heterologous gene expression elicited substantial changes in the cellular transcriptome, while changes in the proteome were of a lesser extent. Transcriptomic changes triggered by plasmid DNA mainly involved the activation of genes related to inflammatory response and antiviral immunity. This cellular response was shared across all experimental conditions used (*i.e*., mock transfections, EGFP expression alone or SHBLE and EGFP expression), pointing towards a common response to the presence of plasmid DNA in the cytoplasm of the transfected cells. Indeed, DNA in eukaryotic cells is restricted to enveloped subcellular structures (nucleus, mitochondria, chloroplasts), and the presence of naked DNA in the cytoplasm is most often related to viral infections. In vertebrates several cytoplasmic DNA sensors exist such as the cyclic GMP-AMP synthase (cGAS) and the stimulator of interferon genes (STING) (**Wu & Chen, 2014**). Consistent with the activation of such cytoplasmic DNA-sensing pathways and their known downstream effects on gene expression (**Stetson & Medzhitov, 2006; Wu & Chen, 2014**), we detect in our transfected cells many activated genes related to Type I interferon signaling pathway. This was the case of upregulation in all our experimental conditions of interferon stimulated genes *ISG15* and *ISG20*, interferon induced proteins with tetratricopeptide repeats *IFIT1, IFIT2* and *IFIT3*, or of the interferon alpha-inducible protein 6 *IFI6* genes. Though our study did not aim at characterizing the HEK293 response to transfection, our results suggest that it induces an inflammatory response that could be dependent on *STING* activation, as shown by a study conducted on this cell line (**Khiar et al, 2017**). Finally, we also observed that transfection led to up-regulation of genes known to have viral RNA sensor roles, such as the four members of the OAS gene family (2’-5’ oligoadenylate synthetases, *OAS1, OAS2, OAS3, OASL*), all of them regulated by interferon signaling and encoding for enzymes displaying RNAse activity (**Kristiansen et al, 2011; Zhu et al, 2014**). Note that this observation is not necessarily in contradiction with the fact that we transfected DNA and not RNA, knowing that crosstalk exists between antiviral sensors of DNA and double-stranded RNA (**Cheng et al, 2007**). Our previous results on this experimental system (**Picard et al, 2022**) further show that transfection alone leads to a 50% drop in cell growth, in line with a functional anti-viral response that would prevent access of exogenous DNA to the cell nucleus upon cell division. In conclusion transfection of plasmid DNA resulted in the induction of a common anti-viral response through the overexpression of mRNAs involved in inflammatory processes, consistent with functional correlates of cell growth decrease.

Expressing plasmid-encoded heterologous genes in our cells triggered large transcriptomic changes. For each of the different synonymous version, we identify at least 480 cellular transcripts to be differentially expressed after transfection. In striking contrast to these transcriptomic changes we observed no differences in the proteomes of the transfected cells. A technical explanation for this differential response between transcriptomic and proteomic responses could be related to the differential power of the technical approaches quantifying transcripts and proteins. In our case we detect the presence of a median 13,661 gene transcripts per sample while we could only detect 2,712 proteins. We also observe that mRNAs for which we detected the corresponding protein were on average 3 times more abundant than mRNAs without protein detected (median log10 TPM values: 1.76 versus 1.28, p < 2.2e-16, Wilcoxon Mann Whitney – Fig. S13). Concomitantly, detected proteins were biased towards those corresponding to transcripts displaying no significant variation of expression after transfection (Fig. S13), providing a potential explanation for the absence of protein differential expression. Despite this technical gap, our results contribute to the growing evidence obtained by combining transcriptomics and proteomics (or Ribo-Seq) approaches, showing that proteomes tend to be more stable, and/or to display a larger inertia to change, than transcriptomes. Such a trend has been communicated for yeasts species (**McManus et al, 2014**), for the molecular responses of other fungi either to stressors or environmental cues (**Blevins et al, 2019; Brancini et al, 2019**), as well as across primates (**Khan et al, 2013**). In this regard, it has been shown that human tissues involved in similar broad functions display similar proteomes, even if their transcriptomes are divergent (**Wang D et al, 2019**) and that this trend is conserved across evolutionary timescales for mammals (**Wang ZY et al, 2020)**. Additionally, differences in degradation rates of mRNAs and proteins can further enhance this gap between the transcriptomic and the proteomic responsiveness: in mammalian cells in culture, proteins display in average five times larger half-lives than mRNAs (46 h *vs*. 9h, respectively), and there is no correlation between the stability of mRNA and the corresponding protein (**Schwanhaüsser et al, 2011**). It is thus conceivable that changes at the proteome level appear buffered because highly expressed and long-lived proteins might still be present at their initial expression level (*i.e*., at the time before transfection occurred) and had not sufficient time to achieve a complete turnover. Nevertheless, even if such effect exists, we anticipate its impact on our experimental data to be limited because we collected our cells for the transcriptomic and proteomic determination 48h after transfection.

It is well known that CUBias strongly influences the expression level of a gene (either directly or indirectly via linked variables such as GC composition, dinucleotide composition, mRNA folding energy or changes in decay pathways – **Chamary & Hurst, 2005; Kudla et al, 2006; Vogel et al, 2010; Boël et al, 2016**; **Cambray, Guimaraes & Arkin, 2018; Courel et al, 2019**. Such *cis*-acting effects of CUBias on gene expression have been thoroughly documented for our *shble-egfp* experimental system (**Picard et al, 2022**). The purpose of the present study was to determine whether and to what extent CUBias may display *trans*-acting effects on the overall cell translation. We specifically explored whether strong expression of a heterologous gene lead to a differential impact on the translation of cellular genes as a function of the match between their CUBias and the CUBias of the overexpressed gene. In mammalian cells, virtually all ribosomes are engaged in translation at any given time (**Princiotta et al, 2003**), and translation is the most expensive step in the biological information flow process (**Lynch & Marinov, 2015**), consuming around 45% of all cellular energy (**Princiotta et al, 2003**). It is thus conceivable that high expression levels of certain genes may come at the expense of a limited expression of other genes, due to competition for non-finite pool of translational resources shared by mRNAs present in the cell. This may be especially the case if highly abundant mRNA species are not efficiently translated (**Shah et al, 2013**) – for example due to a poor CUBias. The existence of such *trans*-effects has been verified in *E. coli* using high-throughput approaches with respect to tRNAs availability (**Frumkin et al, 2018**). Another study on *S. cerevisiae* led to the same conclusion that CUBias of highly expressed genes is key in maintaining the overall translation efficiency of the rest of mRNAs present in the cell (**Qian et al, 2012**). Here we test on a mammalian model the hypothesis that CUBias of highly expressed genes impacts translation efficiency via *trans*-acting effects, linked to competition between heterologous transcripts and cellular transcripts to access tRNAs. We found no evidence of such effects in our experimental system, despite having constructed highly divergent synonymous *shble* versions that are expressed at remarkably high levels. Indeed, heterologous *shble-egfp* transcripts represent more than 1% of transcripts in most samples (*i.e*., above 10,000 TPM) and the SHBLE and EGFP proteins together represent more than 0.6% of the total protein abundance detected for all versions except Shble#6. Results presented in Figure 2 and Figure 3 show no support for the codon usage-specific decrease in translation efficiency of cellular mRNAs expected under the hypothesis that high expression of heterologous genes leads to *trans-*acting effects through tRNA shortage. Regarding the use of rare codons more specifically, which are codons presumably decoded by tRNAs more prone to shortage, the competition hypothesis predicts that overexpression of heterologous genes enriched in rare codons should result in a negative impact on the translation of cellular genes enriched in these same rare codons (**Yona et al, 2013; Frumkin et al, 2018**). Our results however do not support this hypothesis. In contradiction to this expectation, we instead observe a decreased translation efficiency in cellular genes enriched in common codons when we force the cells to overexpress heterologous genes rich in rare codons. We ruled out that genes with decreased translation efficiency are more expressed (Fig. S14), which could have rendered them more prone to potential tRNA shortage independently of their CUBias. We must admit that we have not found yet a satisfying explanation for this counter-intuitive pattern.

Synonymous versions of the *shble* coding sequence used in this study were designed to present distinct CUBias, except for the first seven codons, which are identical across. Thus, the chemical and coding environments immediately surrounding the translation starting point are identical for all synonymous versions and our postulate is that our gene recoding strategy allows to largely tease apart initiation-from elongation-driven effects. Local mRNA structures present around the start codon can hinder ribosomal binding and early progression and hence translation initiation. Indeed, variation in the strength of secondary structures around the start codon is the largest explanatory factor for differences in translation in *E. coli* (**Cambray, Guimaraes & Arkin, 2018**). Molecular modelling suggests indeed that translation is mainly limited by initiation rate rather than by elongation rate (**Shah et al, 2013; Riba et al, 2019**). These predictions are supported by numerous experimental studies reporting a link between mRNA secondary structures immediately upstream the translation start site and protein synthesis. This positive relationship between mRNAs with less structured 5’ UTR and higher protein production seems to be conserved throughout evolution as has been identified in *E. coli* (**Kudla et al, 2009**), unicellular eukaryotic organisms (**Shah et al, 2013; Weinberg et al, 2016; Wang SE et al, 2020**) and mammals including humans (**Gandin et al, 2008; Mauger et al, 2019** but see **De Sousa Abreu et al, 2009** who described no effect of the initiation rate on translation efficiency in human transcripts). Our design for synonymous versions recoding of the *shble* gene focused on the elongation step of protein synthesis, so we don’t expect mRNAs species of the different *shble* versions to be initiated at different rates. Results present in this study and our previous one together suggest that elongation can alter protein expression in *cis-*, but that *trans-*effects on cellular mRNAs translation efficiency are modest and, importantly, not related to their CUBias.

Results presented in Figure 4 suggest that differences in GC composition between our synonymous versions impact host translation efficiency in opposite ways: cultures transfected with AT-rich versions (Shble#3, Shble#4, Shble#6) presented an enhanced translation efficiency of cellular mRNAs rich in GC-ending codons whereas cultures transfected with GC-rich versions (Shble#1, Shble#2, Shble#5) presented an enhanced translation efficiency of cellular mRNAs rich in AT-ending codons. Considering the hypothesis of *trans-* effects exerted on cellular genes through translation of heterologous genes, we expected our synonymous versions to cluster not according to their GC-content but rather according to their overall match with the CUBias of the host cell (proxied here by their COUSIN score). We first note that we did not recover the expected pattern of AT-rich versions triggering the expression of *schlafen11*, which could have explained why translation of GC-rich cellular transcripts seems favored in sample expressing these versions. In mammals, this gene is indeed known to be a part of the innate immune response against viral infections resulting in an altered translation in a CUBias-dependent manner, specifically impairing AT-rich transcripts translation (**Li M et al, 2012**). As a second line of thought, a highly expressed heterologous gene could globally modify the cellular translation dynamics because of the overload it imposes on the translation machinery, via ribosomal sequestration, for instance (**Princiotta et al, 2003**). It has been indeed shown that adenine-rich mRNAs promoted ribosomal binding to these transcripts (**Castillo-Méndez et al, 2012**).

Finally, we would like to draw attention to the fact that modifying CUBias so that the designed sequences are either enriched or depleted in the most frequent synonymous codons found in a genome has not always straightforward consequences on translation efficiency. Albeit it is probably true that in most cases such strategy of codon “optimization” (or “de-optimization”) gives the expected results in terms of protein production (see **Quax et al, 2015** for a review; **Welch et al, 2009; Schmitz & Zhang, 2021**), there are several instances that produced surprising results, not only in humans but also for micro-organisms (**Pop et al, 2014**). In bacteria for example some studies reported that varying CUBias did not change protein expression of the target, was it a transgene or a gene in the genome of a recoded strain (**Kudla et al, 2009; Frumkin et al, 2018**) or worse reported that encoding a protein with putatively most optimal codons ultimately led to a decrease in expression (**Agashe et al, 2013**) or activity (**Zucchelli et al, 2017**). Regarding mammalian cells, a systematic examination of sequence features that impact protein concentration performed on a human cell line revealed for example that CUBias only has a minor impact compared to other sequence features (**Vogel et al, 2010**). A work performed by **Cambray, Guimaraes & Arkin, 2018** dissected out the individual and combined impact of several mRNA compositional variables onto translation (*e.g*. ability of sequences at the 5’-untranslated region at recruiting ribosome, strength of mRNA structures around the translation start, and CUBias) and found the experimental design could only account for hardly 30% of variation in protein synthesis. In conclusion, the last decades have produced a large body of literature about the origin and meaning (if any) of CUBias and its interplay on gene expression, but we find still ourselves facing a puzzle, for which we do not know whether there is actually a solution, especially in the case of humans.

## Materials and Methods

### Design of the synonymous six synonymous plasmidic constructs

The original sequence of the *shble* gene found in *Streptoalloteichus hindustans* was obtained from the GenBank database (X52869.1, https://www.ncbi.nlm.nih.gov/nuccore/X52869.1?report=genbank). Six synonymous coding sequences (CDS) of this gene were designed according to the ‘one codon by amino acid’ rule. Codon usage of these six synonymous versions was designed as follows: version Shble#1 uses exclusively the human most frequent codon; versions Shble#2 and Shble#3 use codons with the highest GC or the highest AT contents among the two most common codons, respectively; version Shble#4 uses exclusively the human least frequent codon; versions Shble#5 and Shble#6 use codons with the highest GC or the highest AT contents among the two least common codons, respectively. An invariable AU1 tag (MDTYRYI) was added at the start of each synonymous version, resulting in the same first seven codons for all synonymous versions. A PCA illustrating how these synonymous versions differ in terms of their CUBias among one another is given in Fig. 1A. Each of these synonymous versions of the *shble* CDS was linked to the CDS of the Enhanced Green Fluorescent Protein (*egfp*) via a P2A self-cleaving peptide sequence, so that a bicistronic mRNA is produced and translated into two proteins after cleavage of the P2A peptide. Synonymous versions were synthesized (Genescript) and cloned into the XhoI site of the pcDNA3.1-P2A-EGFP-C plasmid, that also contains in its backbone a Neomycin resistance gene (*3’-glycosyl phosphotransferase*). In addition to the six constructs bearing different synonymous versions of *shble* translationally fused to *egfp*, a control vector missing the *shble* gene upstream the *p2a-egfp* region was designed, named the Empty version. This Empty version serves as a control to account for *egfp* expression. Note that the sequence encoding *egfp* is similar for all constructs and that its CUBias is strongly biased towards human’s most favored codons, resembling to the CUBias of versions Shble#1 and Shble#2.

### Measurement of codon usage bias

COUSIN scores (Codon Usage Similarity Index, **Bourret, Alizon & Bravo, 2019**) were used to compare the codon usage of both human and plasmidic genes to the average *Homo sapiens* codon usage (https://cousin.ird.fr). The COUSIN score reflects the extent to which codon usage of a query sequence matches the one of a reference. In our study, the reference is the whole codon composition of the coding part of the human genome. Briefly values above 1 reflect an overmatch compared to the codon usage preferences of the reference while values below zero reflect preferences opposite to those of the reference. By design, all six synonymous versions of the *shble* CDS have a different COUSIN values, with some being over-humanized (higher COUSIN; Shble#1 and Shble#2) while others being enriched in infrequent codons compared to the average human codon usage (lower COUSIN; Shble#3 to Shble#6). Ordered by decreasing COUSIN scores, the COUSIN score of the six different synonymous versions based on their CDS are as followed: 3.47 (Shble#1), 3.42 (Shble#2), 0.19 (Shble#5), −0.48 (Shble#3), −1.32 (Shble#6) and −2.53 (Shble#4). The COUSIN score of the *egfp* CDS is high (3.38), close to the one of Shble#1 and Shble#2 versions.

### Cell transfection and sampling design

HEK293 cells (Human Embryonic Kidney cells, CRL-1573, ATCC) were cultured at 37°C and 5% CO2, in Minimum Essential Medium (Earle, M1MEM10K, Eurobio), with 10% FBS (Fetal Bovine Serum, CVFSVF0001, EuroBio) and 1% Penicillin-Streptomycin (15140122, Fisher scientific). Transfection was done in six-well plates, with 1.17×10^5^ cells/mL. The next day, medium was replaced by MEM 2% FBS. Each synonymous plasmid mixed with turbofect reagent (12331863, Fisher scientific) was added in each well and cells were sampled at day 2 (Trypsin-EDTA, CEZTDA000U, Eurobio) for further processing. In total, three independent series of transfection experiments – each with the eight conditions in duplicates – were performed. This resulted in 48 samples, balanced as follows: six Shble#1, six Shble#2, six Shble#3, six Shble#4, six Shble#5, six Shble#6, as well as six Empty samples and six Mock samples.

### Sequencing and RNAseq data analyses

RNA extraction was performed using the Monarch Total RNA miniprep kit (T2010S, NEB), following the manufacturer’s recommendations. Total RNAs were sent to Genewiz NGS laboratory (New Jersey, USA), where they performed polyA selection, strand-specific RNA library preparation and 2×150bp sequencing on an Illumina HiSeq4000 system. Quality checks of raw reads were performed with FastQC (available online at: http://www.bioinformatics.babraham.ac.uk/projects/fastqc). Reads were passed to Cutadapt (v1.10, **Martin, 2011**) to remove universal Illumina adapters then trimmed with Trimmomatic (v0.38, **Bolger, Lohse & Usadel, 2014**) using the following options: PE HEADCROP:13 SLIDINGWINDOW:4:30 MINLEN:85. Processed reads were pseudo-mapped onto the human transcriptome (//ftp.ncbi.nlm.nih.gov/genomes/all/GCF/000/001/405/GCF_000001405.39_GRCh38.p13, 19,812 predicted protein coding genes). To allow reads coming from expression of the genes contained in the plasmid to be aligned, the transcriptome was appended with the Neomycin resistance transcript sequence (*Neo-R*) and with the *shble#X-p2a-egfp* transcript sequence (where X stands for 1 to 6; *i.e*., for each sample transfected with Shble#1 to Shble#6, the corresponding synonymous sequence of shble was used to allow proper mapping). For Empty, only the *egfp* transcript sequence was added. For Shble#4 and Shble#6 samples, spliced forms of the shble-p2a-egfp transcript identified in a previous study (**Picard et al, 2022**) were also added to allows reads spanning the junctions to be aligned. Pseudomapping onto the transcriptome was performed with Kallisto (v0.46.0, **Bray et al, 2016**) with default options except for the number of bootstrap samples and the number of threads, that were respectively set as follows: -b 100 -t 16. As outputs of Kallisto, we retrieved quantification at the gene level in raw read counts and in Transcripts Per Million (TPM), both for cellular and heterologous transcripts.

### Cellular transcripts level distribution over the 48 transcriptomes

We plotted the distribution of expression of all cellular transcripts for each transcriptome using log10(TPM+1) values and reported the number of transcripts expressed above 1 TPM (Fig. S15). We calculated the correlation of transcriptome-wide expression between duplicates derived from the same transfection experiment. Extremely high R2 values obtained for such pairs of duplicates (in all cases R2 > 0.99 – Fig S16) led us to average TPM obtained from the two repetitions of each transfection, thereby reducing the number of samples from 48 to 24. For all remaining analyses we used these “collapsed” measurements (*i.e*., n=3 independent quantification per condition).

### Differential expression of cellular transcripts after 2-days post transfection by different synonymous constructs

After duplicates collapsing, our transcriptomic dataset was reduced from 48 to 24 data points, with each condition supported by n=3 independent experiments. We defined for each condition the set of expressed mRNAs as those expressed above 1 TPM in at least two out the three replicates that correspond to the condition. On average across all eight conditions 13,650 mRNAs were detected as being expressed (min = 13,297; max = 13,863). Working on the union of these sets of expressed mRNAs (14,139 mRNAs), fold-changes relative to the Mock condition were calculated for each seven conditions (Shble#1 to Shble#6 + Empty). Genes were considered differentially expressed relatively to the Mock in a condition if the median fold-change (across the three samples corresponding to this condition) was above 2 or below 0.5 for this gene. In parallel, we also used the DESeq2 R package (**Love, Huber & Anders, 2014**) on the total set of 19,812 mRNAs to similarly identify differentially expressed (DiffExp) transcripts in comparison to the Mock after transfection by our different constructs. For this, we used counts estimated by Kallisto, that we normalized using the *estimateSizeFactors(*) function of DESeq2. This second approach presents the advantage of returning corrected p-values and takes into account the lack of power of differential expression for genes with low counts. Overlaps of genes identified as differentially expressed by the two distinct approaches were considered for the analysis of DiffExp transcripts.

### Protein extraction and label-free proteomic analysis

For protein extraction, the two duplicates sampled from the same transfection experiment were pooled in one tube, thereby reducing the number of samples from 48 to 24. Solubilized proteins were resuspended in Laemmli buffer and 20-30 μg of proteins were stacked on a SDS-PAGE gel. Proteins were in-gel digested using Trypsin (Trypsin Gold, Promega), as previous described in **Shevchenko et al, 2006**. Proteomic data were collected in data dependent acquisition mode using a Q Exactive HF mass spectrometer coupled with Ultimate 3000 RSLC (Thermo Fischer Scientific). The software MaxQuant (**Cox & Mann, 2008**) was used to analyze tandem mass spectrometry data. All m/Z spectra were searched using standard settings with the search engine Andromeda (**Cox et al, 2011**) against a target decoy database to deliver false-positive estimations. The database contains entries from the *Homo sapiens* Reference Proteome (UP000005640, release 2019_02, https://www.uniprot.org) and a list of potential contaminants. Sequences of the two heterologous proteins of interest (SHBLE and EGFP) were added into this database to allow for their identification. Search parameters were let at their default values, oxidation (Met) and acetylation (Nt) were applied as variable modifications and carbamidomethyl (Cys) as a fixed modification. False Discovery Rates (FDR) of peptides and proteins identification were let at their default values (both at 1%). Proteins groups were automatically constructed by MaxQuant. A representative ID in each protein group was automatically selected using an in-house bioinformatics tool. After excluding usual contaminants, 4,005 proteins were identified in at least one sample (out of 24 samples), of which 2,891 were detected in all the three replicates of at least one condition (Table S3). Protein quantifications in intensity-Based Absolute Quantification (iBAQ, **Schwanhäusser et al, 2011**) were retrieved and normalized by the total sum of iBAQ within each sample (heterologous proteins included) to obtain relative iBAQ (riBAQ), a measure representing the molar fraction of each protein for within-sample normalization (**Shin et al, 2013**). The distribution of riBAQ for proteins detected in each sample is given as log2 scale in Fig. S5. Correlation of riBAQ values between replicates was checked (Fig. S17). Protein quantification in Label Free Quantification (LFQ) was also retrieved for the analysis of differentially expressed proteins (**Cox et al, 2014**). Details regarding protein expression in the different conditions are given in Table S3.

### Analysis of differentially expressed proteins

The software Perseus (**Tyanova et al, 2016**) was used to identify human proteins which level of expression vary depending on the version of the plasmid that was transfected into our cells. Adding the Mock condition in addition to the six synonymous versions and the Empty construct, this resulted to a total of eight groups, each supported by n=3 biological replicates. We used LFQ values for this analysis of DiffExp proteins as this metrics is tailored for comparisons of the expression level of a given protein across different samples. LFQ values were log2 transformed and the following rationale was applied: proteins that were not present in at least 2 out of 3 replicates in a least one condition were filtered out. Only 1,989 proteins with a maximum of one missing log2(LFQ) value were hence retained for the rest of the analysis as they probably represent proteins identified with sufficient level of confidence in at least one condition. Note that this number is low due to the inherent calculation of LFQ values, which returns zero when there is not enough information for a peptide to be detected (**Cox et al, 2014**).

### Analysis of proteins with a qualitative expression pattern (ON/OFF) according to the different conditions

Complementary to the analysis of differentially expressed proteins described above, we selected proteins that displayed qualitative differences (i.e., ON/OFF) across our eight groups (Shble#1 to Shble#6, Empty, Mock). To do so, we selected – based on their riBAQ – proteins that were detected in all three replicates of at least one condition but not detected in any of the three replicates in at least another condition. This stringent definition of qualitative expression pattern yielded 369 proteins displaying such behaviour. Within each of the 24 samples, we found that those expressed in the considered sample among this set of 369 proteins systematically displayed lower riBAQ values than the rest of the proteins expressed in that sample (Fig. S18).

### Calculation of protein-over-mRNA ratios for samples expressing over-humanized versions compared to ratios in the Mock condition

We tested for competition between translation of mRNAs from plasmidic genes that use codons close to the human average and translation of host mRNAs. To do so, we calculated separately for Shble#1, Shble#2 and Empty conditions the ratio between the protein-over-mRNA ratio (riBAQ/TPM) of a given gene in the considered condition and in the Mock condition. For each condition, ratios compared to Mock ratios were calculated from the n=3 samples of the condition and the n=3 Mock samples. After excluding genes that were not detected in all the three Mock samples, we retained 2,471 genes for which we calculated these [riBAQ/TPM]_Condition_ / [riBAQ/TPM]_MOCK_ ratio of ratios.

### Changes in individual protein-over-mRNA ratios for different direction of CUBias and levels of heterologous protein expression

Transfected samples were separated depending on the CUBias of the heterologous genes they encoded: nine samples expressing heterologous proteins with an over-humanized CUBias (n=3 Empty samples, n=3 Shble#1 samples, and n=3 Shble#2 samples) and twelve samples expressing *egfp* plus a version of *shble* rich in rare codons (n=3 Shble#3 samples, n=3 Shble#4 samples, n=3 Shble#5 samples, and n=3 Shble#6 samples). Adding the three samples of the Mock condition, this led to two sets of samples (n=12 and n=15) that do not overlap except for the three Mock samples. Note that Mock samples were included to serve as «anchors» for our linear regressions, representing what happens in the absence of heterologous protein expression. Independently for these two mutually exclusive sets of samples, we used linear regressions to model how protein-over-mRNA ratios (riBAQ/TPM) of a given gene varied with expression levels of over-humanized (n=9 + the three Mock) or rare codons rich (n=12 + the three Mock) heterologous proteins (EGFP + SHBLE). The range of the x-axis for these two sets of regression is provided in Fig. S19. Before running regressions, we filtered genes based on their values of riBAQ/TPM ratio in order to calculate a slope only when considered sufficiently meaningful. We used the following rationale: when the regression was performed based on the n=12 samples, we fixed a threshold of at least 8 non-null values of ratio and when the regression was performed based on the n=15 samples we fixed a threshold of at least 9 non-null values of ratio. This results in 2,554 and 2,585 genes to analyze for regressions based on n=12 and n=15 samples respectively, that later became 2,550 and 2,580 after having removed the very few genes with more than three infinite values of ratio. In each case, genes for which the F-test testing the significance of the regression slope had a p-value smaller than alpha = 5% were considered for further analysis.

### Comparison of COUSIN scores distributions between different gene sets

We compared the distribution of the COUSIN scores between sets of genes using Anderson-Darling tests, with the use of the *ad_test(*) function implemented in the R package “twosamples” (https://github.com/cdowd/twosamples).

### Relative synonymous codon frequency (RSCF) of a coding sequence

For a given CDS, the number of occurrences of each 59 amino acid-encoding codons (Tryptophane and Methionine excluded) is retrieved. Acknowledging the genetic code redundancy, occurrence of each synonymous codons is divided by the sum of occurrences of codons encoding the corresponding amino acid to obtain RSCF values. By construction, summing RSCF values of synonymous codons returns 1, and summing all 59 RSCF values returns 18 (*i.e*., all amino acids except Tryptophane and Methionine). Regarding synonymous versions of *shble*, RSCF profiles allowed sequence discrimination based on CUBias (Fig. 1A).

### Assessment of synonymous codon content in sample’s proteomes compared to transcriptomes using the Prot-to-RNA RSCF metrics

For every annotated gene in the human genome the number of occurrences of each 59 amino acid-encoding codons (Tryptophane and Methionine excluded) was retrieved using the CDS of its longest predicted transcript. Then using expression levels of either all detected cellular mRNAs, only those with the corresponding protein detected by LC-MS/MS (both measured in TPM) or all detected cellular proteins (measured in riBAQ), we quantified for each codon the amount of its “expression” in the transcriptome or in the proteome of our 24 samples as follows: ∑_*g=Gene*_ *Codoncount_g_ · Expression_g_* (**Gingold, Dahan & Pilpel, 2012; Schmitt et al, 2014**) Codon content in each sample was hence represented by two vectors of length 59, one derived from its transcriptome profile of expression and another derived from its proteome profile of expression. From these vectors we obtained vectors of transcriptome and proteome relative synonymous codon frequency (RSCF), by dividing values of each codon encoding a similar amino acid by the total sum of values of codons encoding the considered amino acid (as described in the above section). Ultimately, to quantify how the CUBias ‘flows’ from transcriptome to proteome we divided the proteome RSCF of each sample by its transcriptome RCSF counterpart. We called this metrics Prot-to-RNA RSCF and considered it as a proxy for whole-cell translation efficiency of synonymous codons. To compute this Prot-to-RNA RSCF metrics, we used exclusively transcripts expression data from transcripts that matched protein detected by LC-MS/MS (*i.e*., we did not consider whole-transcriptomes). Note that our codon counts vectors were computed without considering heterologous genes in the transcriptome and proteome.

### Content specificity of sets of differentially expressed (DiffExp) genes using combinatorial configurations

We used a combinatorial approach to estimate to what extent the content of the six sets (Shble#1 to Shble#6) of DiffExp cellular transcripts compared to the Mock was version-specific or redundant across versions. For this, for a given number N of sets included (N from 1 to 6), we calculated the number of unique genes present in the union of these N sets. For each collection of size N, we reported the median number of unique genes obtained from all possible arrangements. The number of possible arrangements of size N drawn among six sets is 6, 15, 20, 15, 6, 1, for N ranging from 1 to 6, respectively.

### Regression of Principal Component scores against *shble* sequence features by accounting for heterologous expression

To account for expression of heterologous transcripts when regressing Prot-to-RNA RSCF PC scores against *shble* sequence features (COUSIN score and GC-richness), we performed a linear regression of Prot-to-RNA RSCF PC scores against *shble-egfp* transcript expression, retrieved the residuals and performed a second linear regression using these residuals as response variable against *shble* sequence features.

### Functional enrichment analysis

The Functional Annotation Chart module of the DAVID tool (https://david.ncifcrf.gov/, **Dennis et al, 2003**) was used to detect functional categories over-represented in gene sets. Functional categories displaying a Fold-Enrichment above 2 and a FDR-corrected p-value below 0.05 were considered as significantly over-represented.

### Statistical analyses

All statistical tests were performed with R (version 3.6.3). Principal components analyses were performed using the *prcomp(*) function of the R package “stats”. Principal components (the eigenvectors of the covariance matrix) were visualized and superimposed to PCA graphs using prcomp()$rotation. Except when explicitly noted, all reported correlation coefficients correspond to Pearson correlation coefficients.

## List of supplementary tables

**Table_S1.xlsx:** List of differentially expressed mRNAs in each condition of transfection versus the Mock condition.

**Table_S2.xlsx:** Significant enrichment in functional categories related to innate and antiviral immunity among DiffExp transcripts and DiffExp transcripts belonging to these categories according to synonymous versions.

**Table_S3.xlsx:** Sets of proteins detected in LC-MS/MS depending on different criteria

**Table_S4.xlsx:** Intra-sample covariation between proteins and mRNAs levels

**Table_S5.xlsx:** Categories of genes according to their variation in riBAQ/TPM ratios with varying heterologous protein expression across conditions

**Table_S6.xlsx:** Aggregated counts of codons in the different samples, obtained by ponderating codon occurrence in transcripts and proteins by their expression levels

## List of supplementary figures

All supplementary figures are grouped in a single pdf file

