## Supplementary Figures for "Human cellular homeostasis buffers *trans*-acting translational effects of heterologous gene expression with very different codon usage bias"

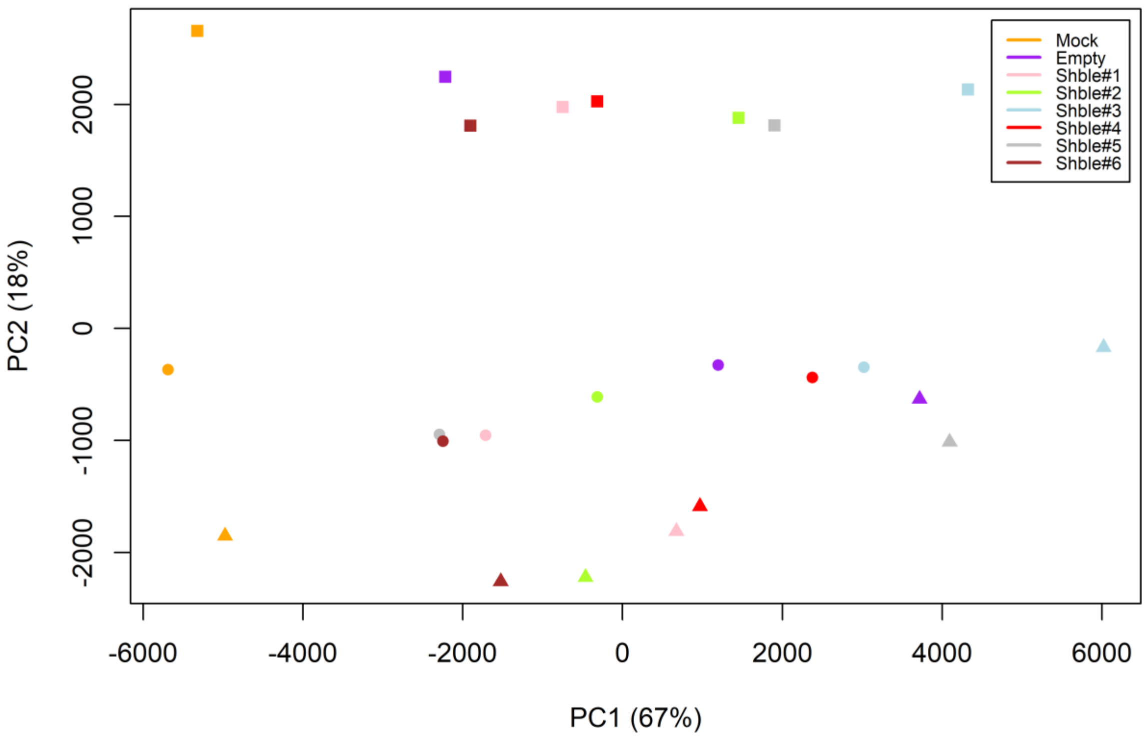


**Fig. S1. Transcriptomic variation across samples.** The first two axes of the Principal Component Analysis performed for the transcriptomes, using exclusively cellular transcripts (*i.e.*, heterologous transcripts excluded) are shown. Values in parentheses represent the fraction of the total variance captured by the corresponding axis. Samples are color-coded according to the transfected construct as in the manuscript: Shble#1 in pink, Shble#2 in green, Shble#3 in light blue, Shble#4 in red, Shble#5 in grey, Shble#6 in brown, EMPTY in purple and Mock in orange. The three experimental batches are symbolized by different shapes.

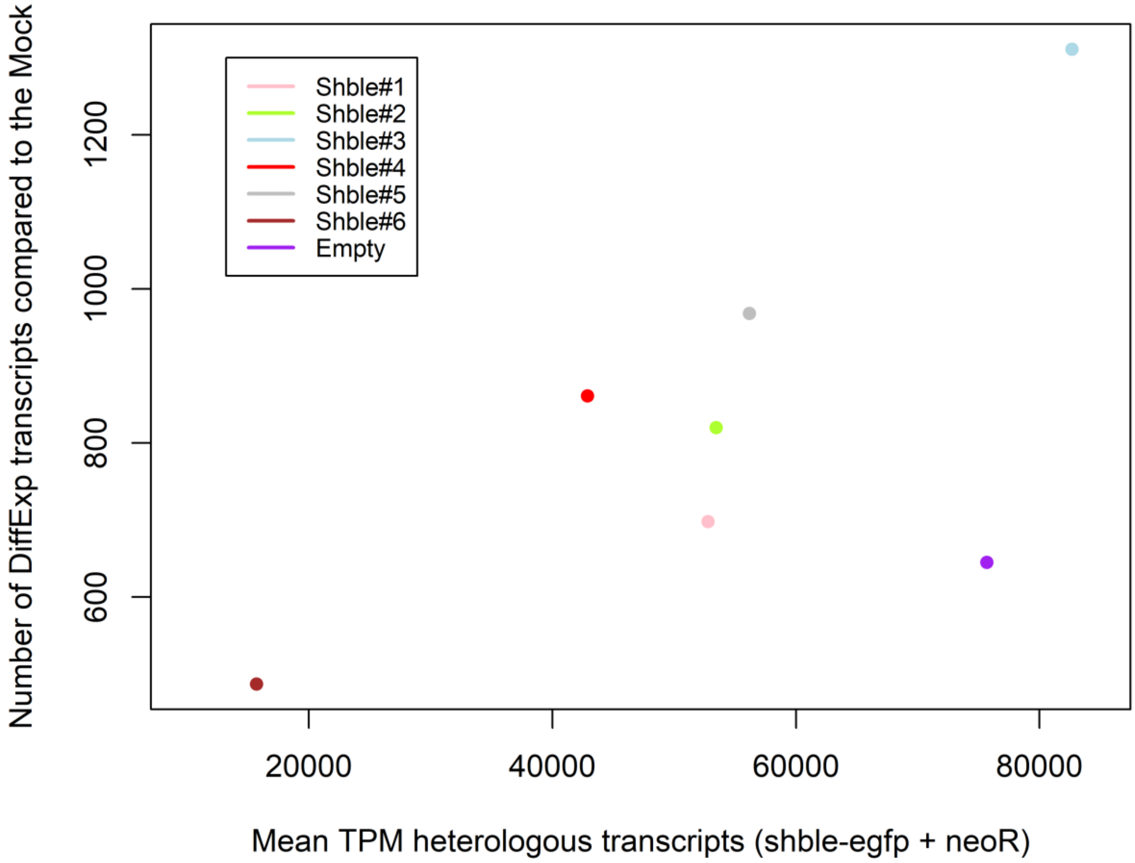


**Fig. S2. Link between the amount of differentially expressed cellular transcripts relatively to the Mock and the mean quantity of heterologous transcripts produced.** For each condition the mean expression of heterologous mRNAs (*shble-egfp* transcript + *neoR* transcript) is the average obtained from the n=3 transfection replicates. Expression levels are given as Transcripts Per Million (TPM). Samples are color-coded as follows: Shble#1 in pink, Shble#2 in green, Shble#3 in light blue, Shble#4 in red, Shble#5 in grey, Shble#6 in brown and EMPTY in purple. The positive relationship observed between DiffExp transcripts and heterologous transcripts expression is consistent with results presented in Figure 1.B of the manuscript.

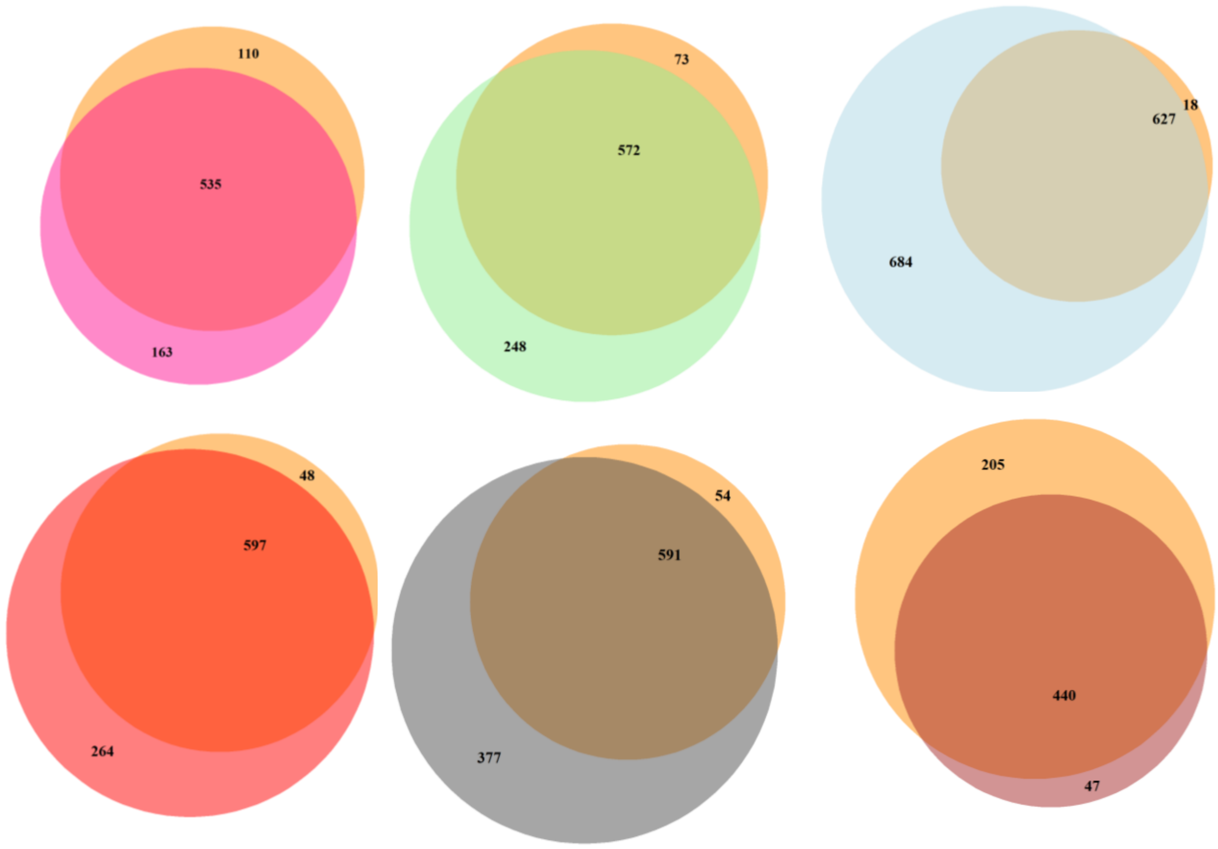


**Fig. S3. Overlap between genes identified as differentially expressed (DiffExp) for each synonymous *shble* version and the set of 645 (DiffExp) genes in the EMPTY condition.** For each version, the proportion of DiffExp genes (relatively to the Mock) that are also DiffExp in the EMPTY condition (salmon background) is as follows: 77% for Shble#1 (pink); 70% for Shble#2 (green); 48% for Shble#3 (light blue), 70% for Shble#4 (red), 61% for Shble#5 (grey) and 90% for Shble#6 (brown).

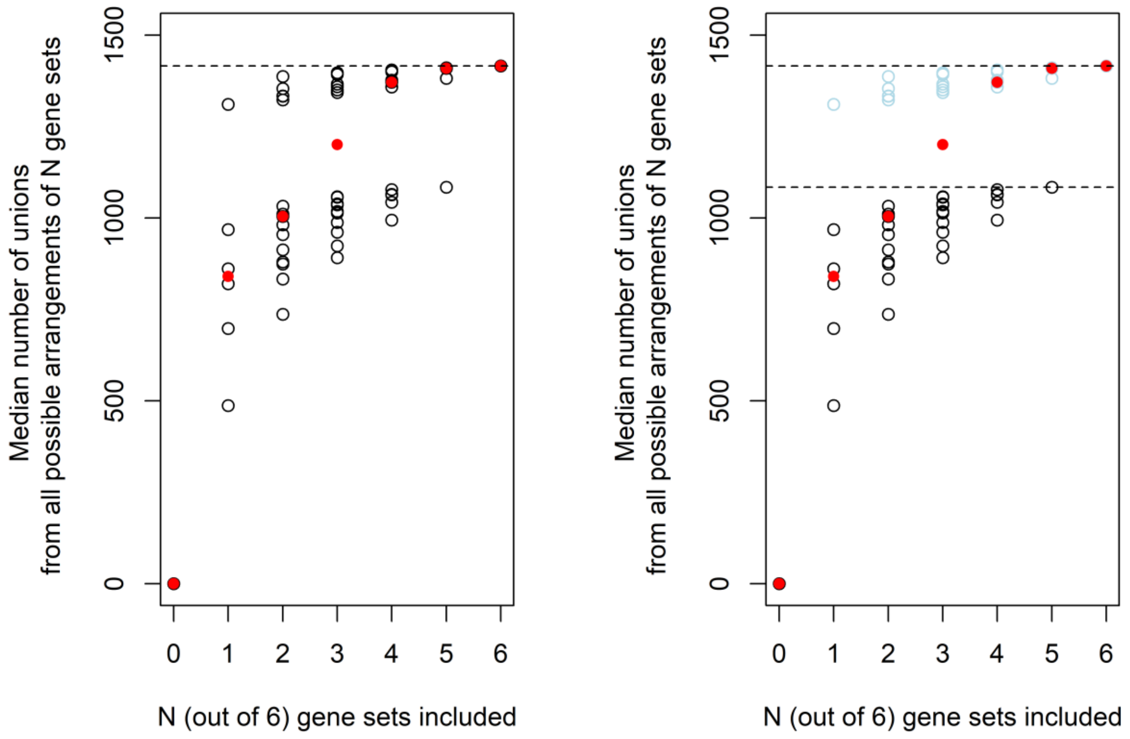


**Fig. S4. Number of differentially expressed (DiffExp) transcripts identified in cells transfected with the different synonymous versions.** *Left*: Values along the x-axis represent how many out of the six sets of DiffExp transcripts (corresponding to conditions Shble#1 to Shble#6) were included, and values along the y-axis represent the number of unique DiffExp genes obtained across these sets. The graph should be read as follows, using the value x=3: for each of the twenty different combinations of three data sets sampled among the six *shble-egfp* conditions, the number of unique DiffExp transcripts are shown in open black circles, while the median of these values is plotted as a red dot (median value of y=1200.5 across the 20 combinations in this example). The horizontal dash line corresponds to the full universe/union of the 1,425 DiffExp genes identified when all six sets of DiffExp genes are included and represents the upper limit of genes that can be identified with sub-sampling N out of the six genes sets. Hence, by combining only half of the six conditions (x=3), a large proportion (85%) of unique genes detected when all six conditions are included is already recapitulated. *Right*: Combinations of gene sets that include the set DiffExp genes in the Shble#3 condition are shown in light blue.

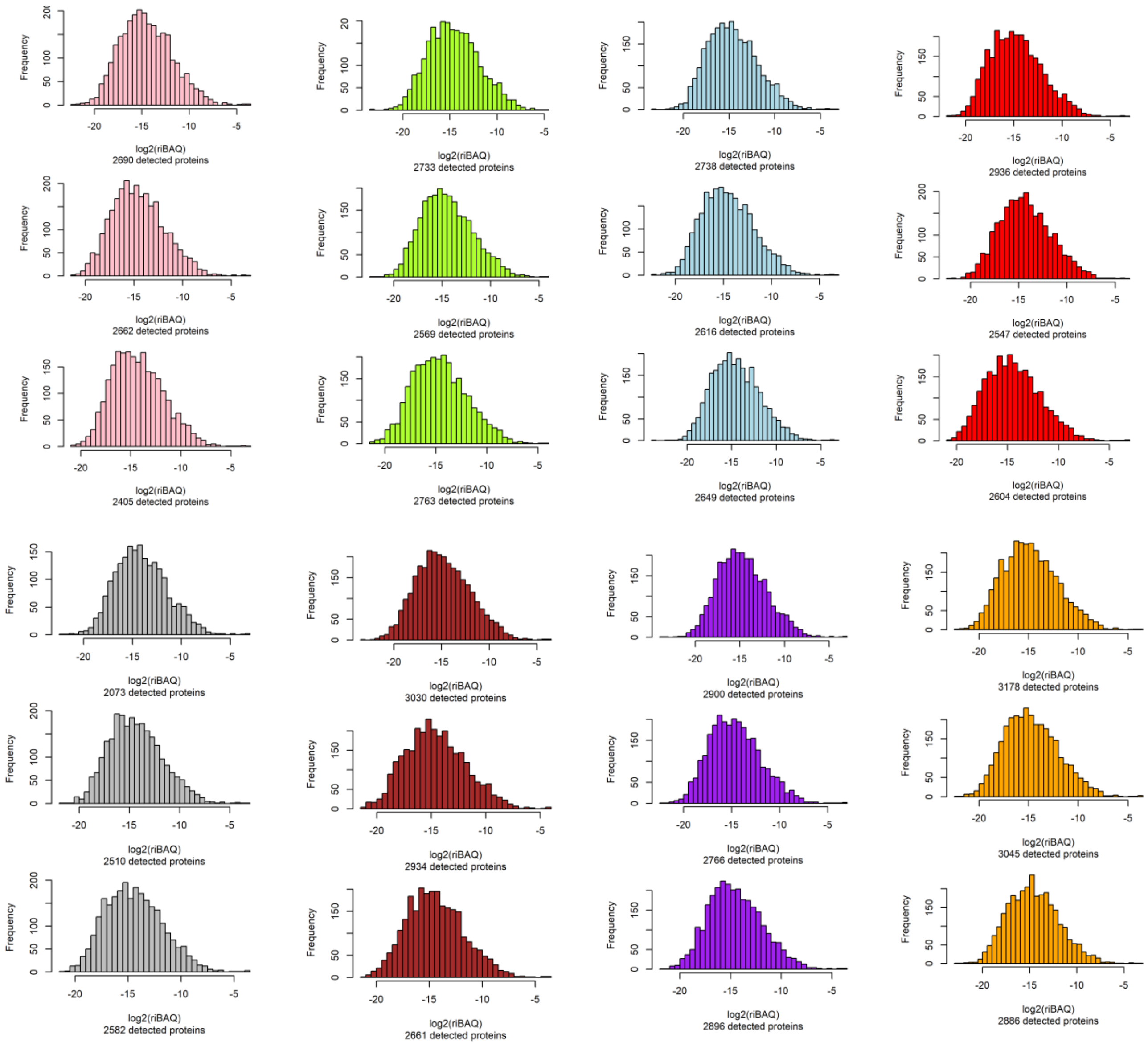


**Fig. S5. riBAQ intensity of detected proteins per sample.** For each condition, the riBAQ distribution in log2 scale is shown for the three independent replicates, with the total number of protein entries detected indicated below the graph. Colors are as follows: Shble#1 in pink, Shble#2 in green, Shble#3 in light blue, Shble#4 in red, Shble#5 in grey, Shble#6 in brown, EMPTY in purple and Mock in orange.

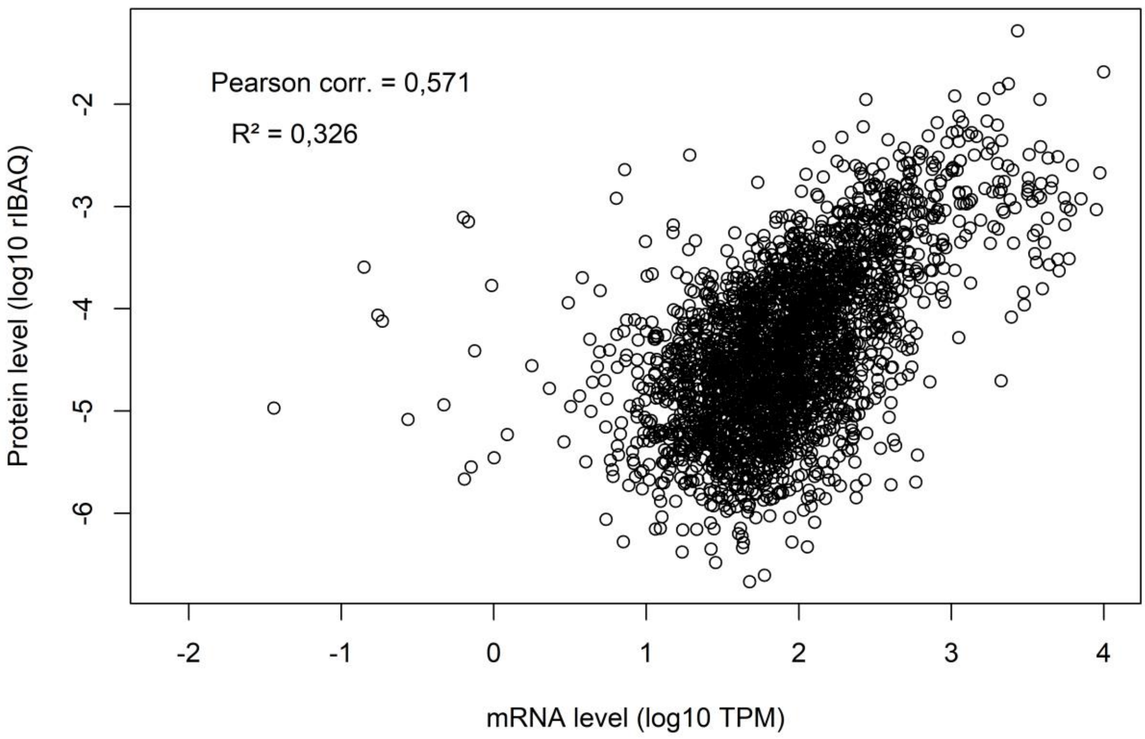


**Fig. S6. Intra-sample covariation between mRNAs and proteins levels.** A representative example is shown, using one sample from the Shble#1 condition. This covariation degree between mRNAs and proteins levels was largely similar to this one for all other samples (Table S4 for details).

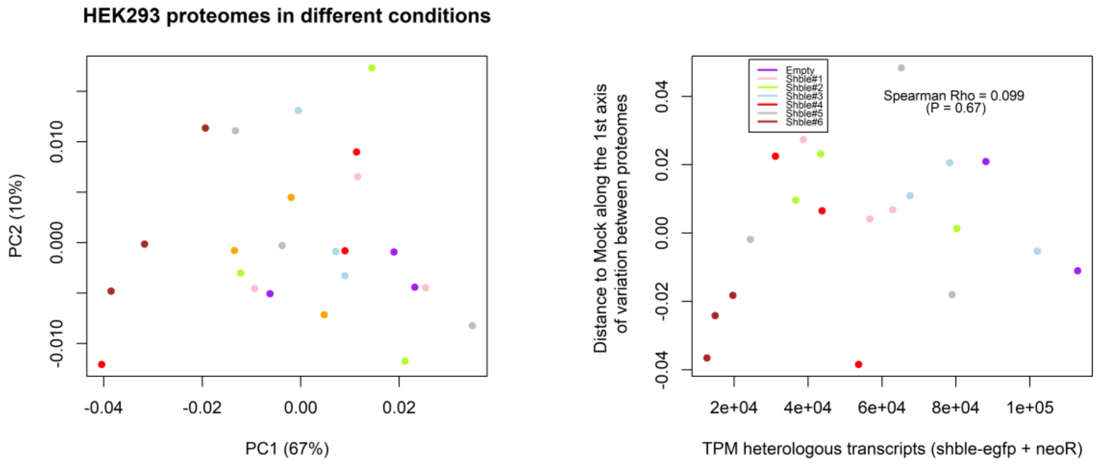


**Fig. S7. Proteomic variation across samples.** Left: The first two axes of the Principal Component Analysis performed for the proteomes (expressed as relative intensity-Based Absolute Quantification – riBAQ – values), using exclusively cellular proteins (*i.e.*, heterologous proteins excluded) are shown. Values in parentheses represent the fraction of the total variance captured by the corresponding axis. Right: the distance between transfected and Mock samples, obtained as the difference between their projections onto the first axis of the PCA is plotted against heterologous transcripts expression levels (see Figure 1.B of the manuscript for a similar analysis using transcriptome data). Samples are colour-coded according to the transfected construct as in the manuscript.

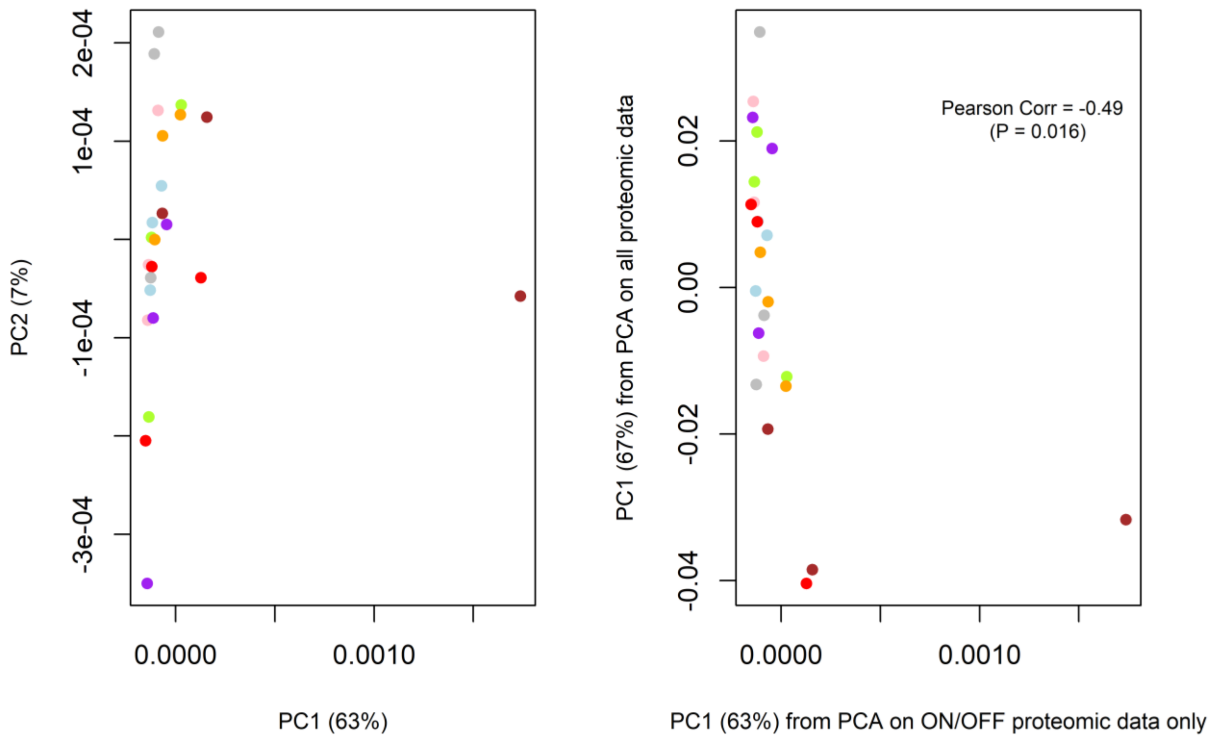


**Fig. S8. Variation across samples based on proteins displaying a qualitative pattern of expression (ON/OFF) and comparison with the profile obtained when all proteomic data are considered.** Left: PCA based on riBAQ from cellular proteins expressed in at least one condition but not in another one (see the Methods section of the manuscript and Table S3). Right: Correlation between first axes of variation of the PCA shown in left – that considers only ON/OFF proteins – and the PCA considering all protein expression data shown in Fig. S7.

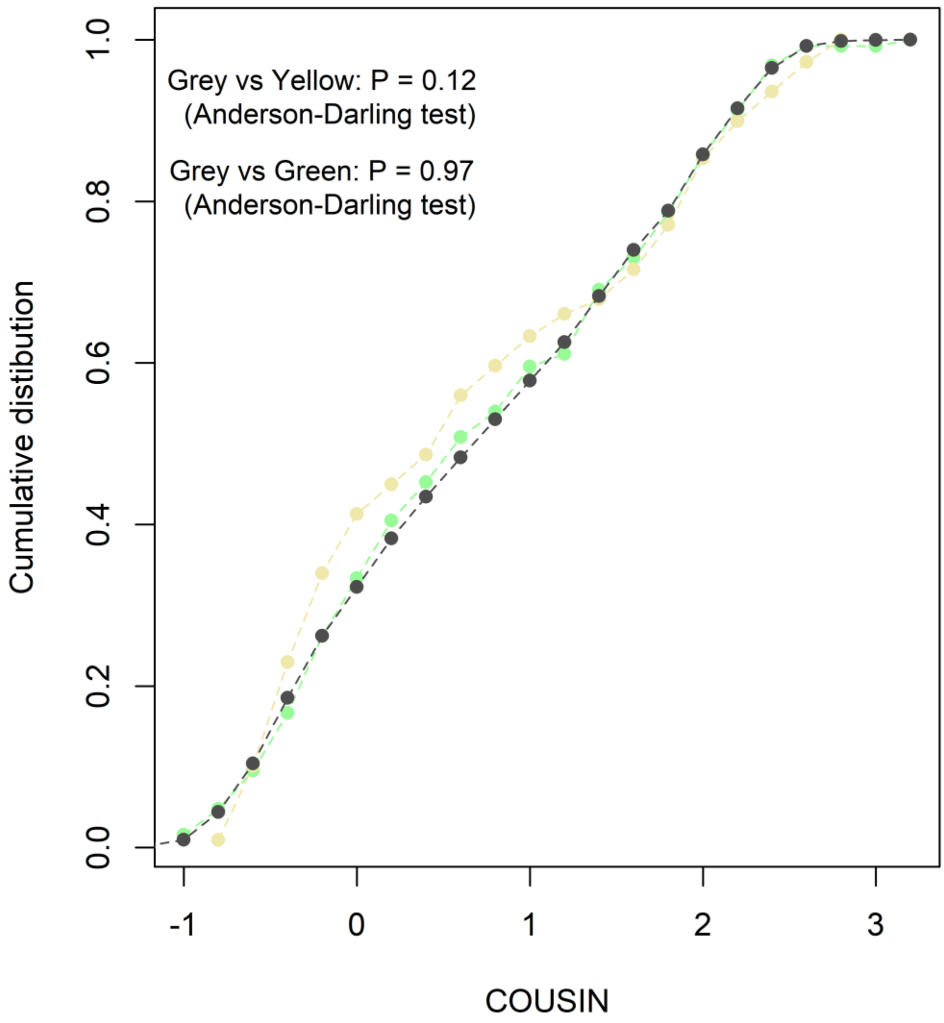


**Fig. S9. Cumulative distribution of COUSIN scores for genes displaying changing riBAQ/TPM ratio values with increasing heterologous protein expression from EMPTY, Shble#1 and Shble#2 samples.** For the set of genes displaying negative changes (n=126) the distribution is shown in green and for the set of genes displaying positive changes (n=109) the distribution is shown in yellow. The underlying cumulative distribution of the 2,550 genes included in this analysis is plotted in grey. P-values of Anderson-Darling tests comparing distributions are indicated. The comparison between the green and yellow cumulative distribution is presented in Fig. 3C of the manuscript.

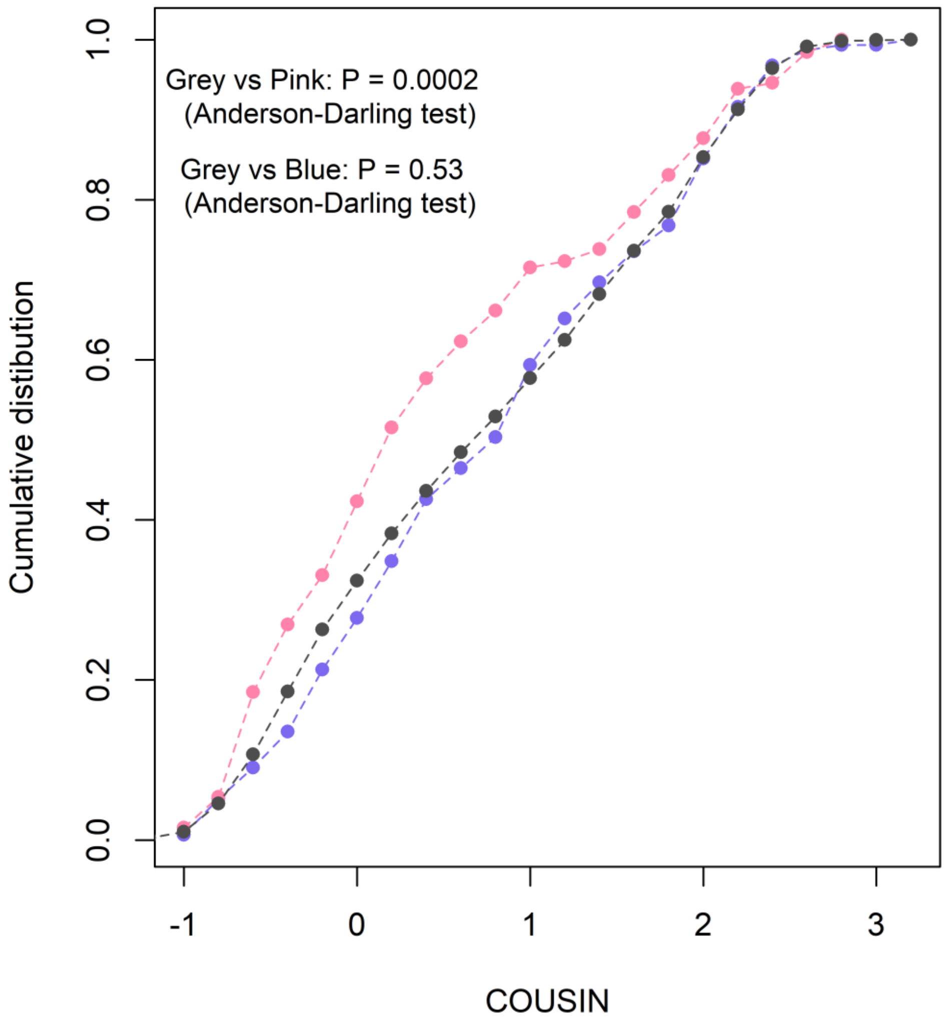


**Fig. S10**. **Cumulative distribution of COUSIN scores for genes displaying changing riBAQ/TPM ratio values with increasing heterologous protein expression Shble#3, Shble#4, Shble#5 and Shble#6 samples.** For the set of genes displaying negative changes (n=155) the distribution is shown in blue and for the set of genes displaying positive changes (n=130) the distribution is plotted in pink. The underlying cumulative distribution of the 2,580 genes included in this analysis is given in grey. P-values of Anderson-Darling tests comparing distributions are indicated. The comparison between the pink and blue cumulative distribution is presented in Fig. 3D of the manuscript.

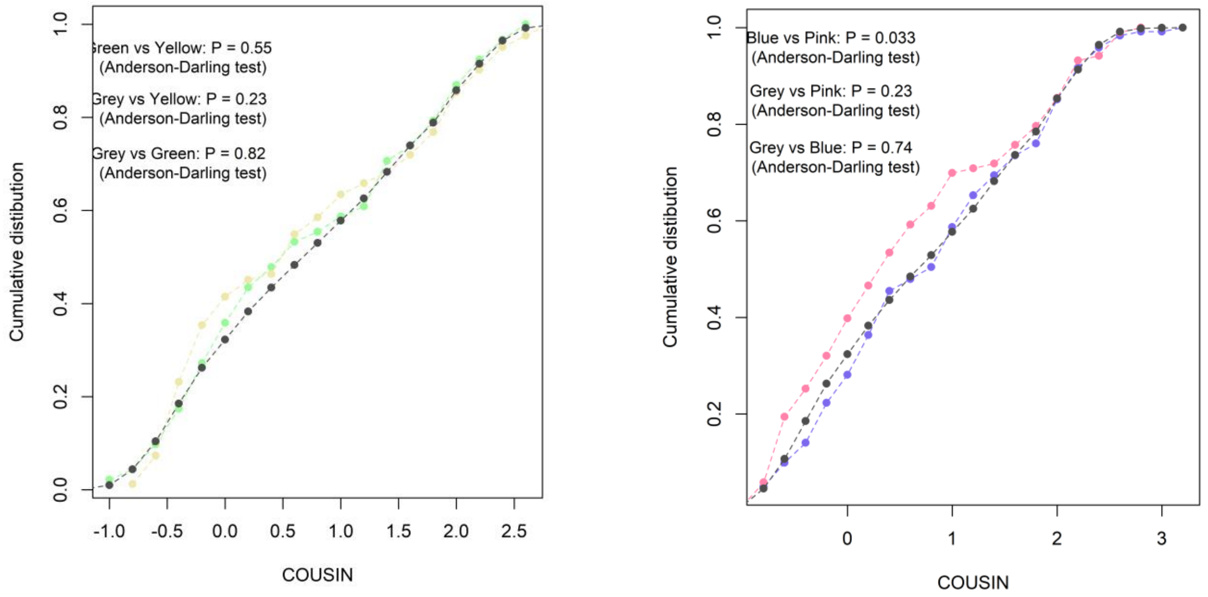


**Fig. S11.** Left: Same as Figure 3C of the manuscript, except that the 27 genes positively impacted by over-humanized protein expression that were also positively impacted by expression of rare-codons enriched versions had been removed (resulting in n= 82 genes, yellow set) and that the 34 genes negatively impacted by over-humanized protein expression that were also negatively impacted by expression of rare-codons enriched versions had been removed (resulting in n=92, green set). Right: Same as Figure 3D of the manuscript, except that the 27 genes positively impacted by rare-codon enriched protein expression that were also positively impacted by expression of over-humanized versions had been removed (resulting in n=103 genes, pink set) and that the 34 genes negatively impacted by rare-codon enriched protein expression that were also negatively impacted by expression of over-humanized versions had been removed (resulting in n=121, blue set).

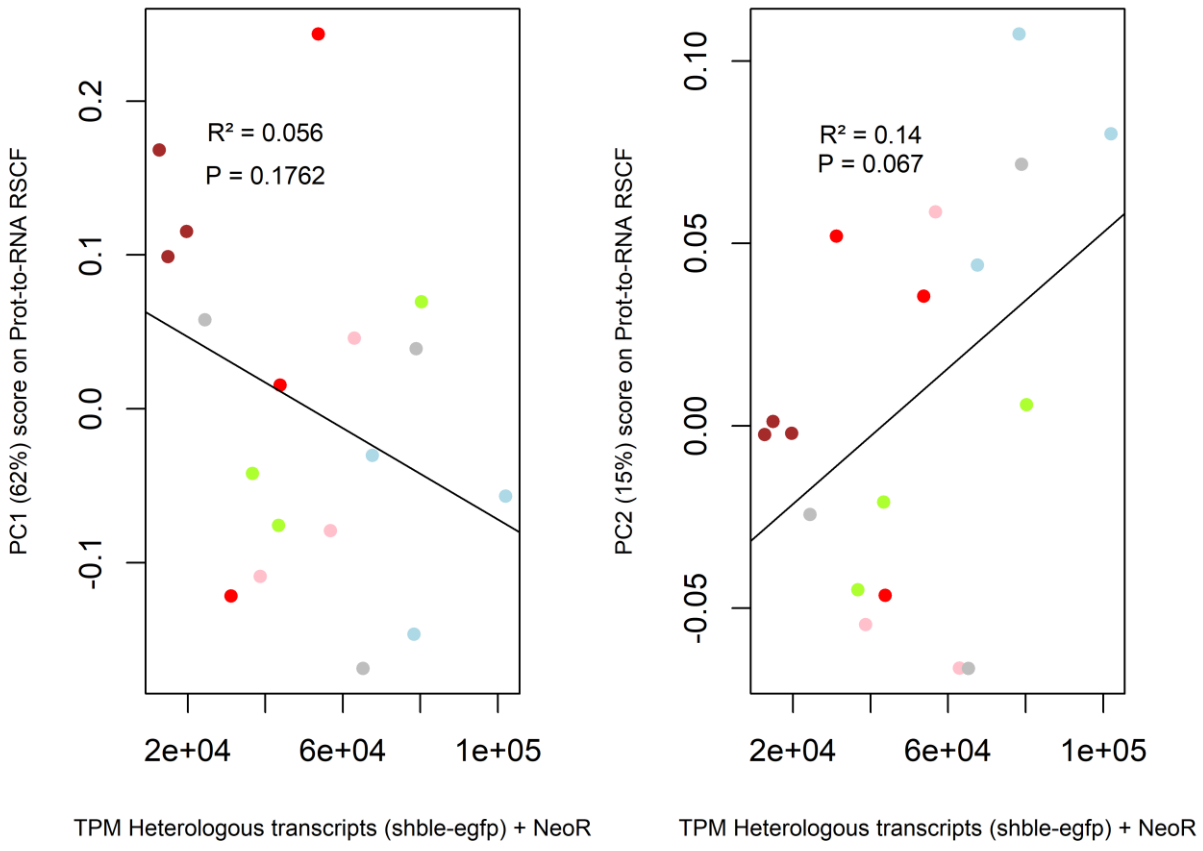


**Fig. S12. Link between Prot-to-mRNA RSCF variation across samples and the quantity of heterologous transcripts expressed.** Scatter plots between the amount of heterologous transcripts and scores along first (left) and second axes (right) of variation across samples for their RSCF values (PCA presented in the Figure 4B of the manuscript). The Pearson correlation coefficient and its associated test of significance are given.

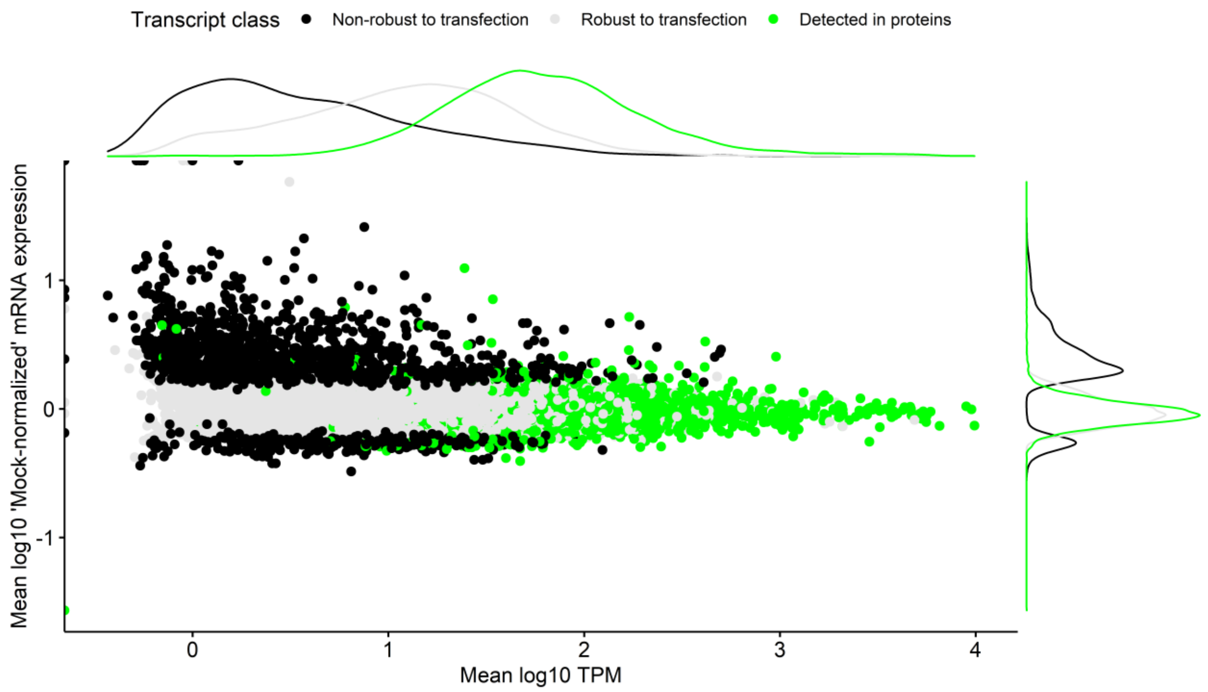


**Fig. S13. Combined patterns of expression level and expression change after transfection for different classes of transcripts.** Transcripts were categorized into three mutually exclusive classes: transcripts with the corresponding protein detected in LC-MS/MS(4,390 transcripts, green), transcripts DiffExp after transfection (*i.e.,* non-robust) in at least one condition (1,425, black), and transcripts robust to transfection and without protein detected (13,997, grey). Genes with the corresponding protein detected (green) were found to be those displaying highly expressed transcripts (x-axis) and whose abundance is not altered by transfection (y-axis).

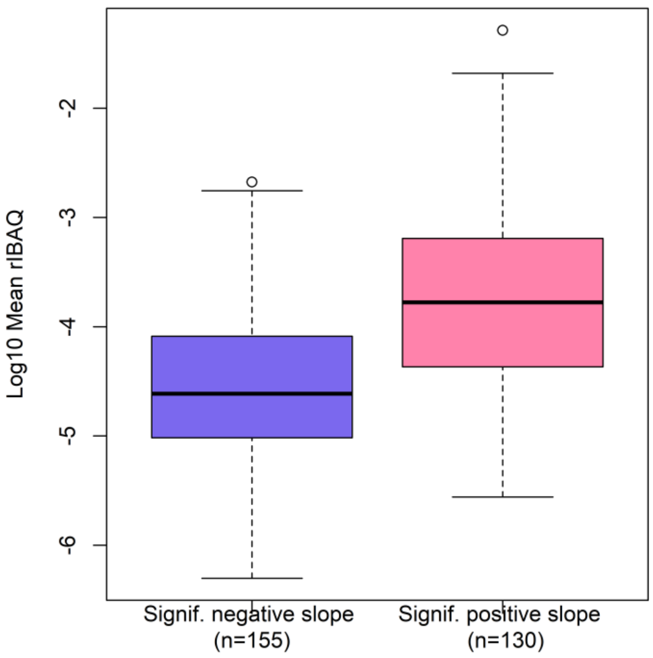


**Fig. S14. Protein levels for genes displaying changing riBAQ/TPM ratio values with increasing heterologous protein expression from Shble#3, Shble#4, Shble#5 and Shble#6 samples.** For the two sets of genes, their expression level averaged across all four conditions is displayed. The difference is significant with a Wilcoxon-Mann-Whitney test ($P < 2.2e-16$).

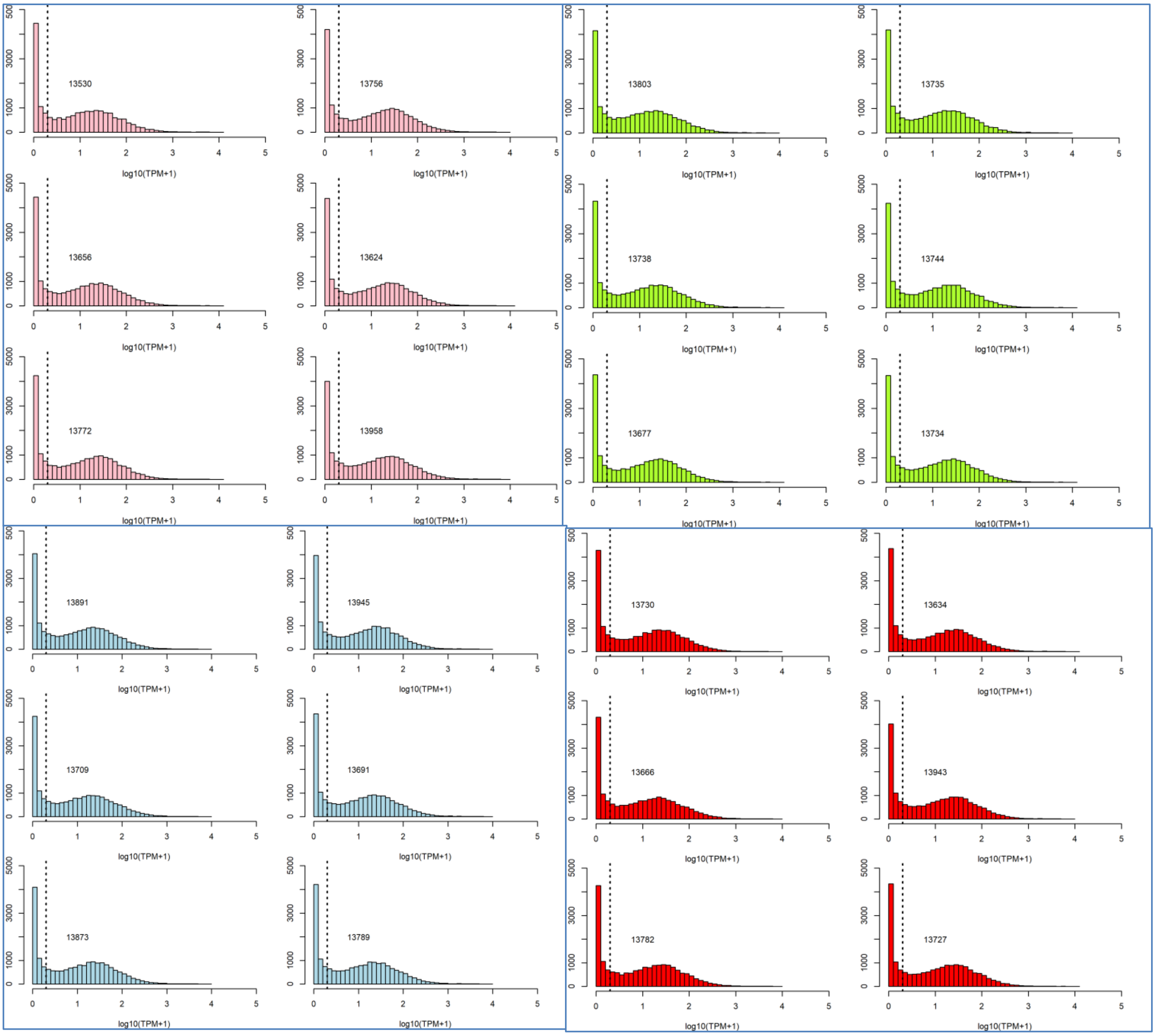
13530
13756
13803
13735
13656
13624
13738
13744
13772
13958
13677
13734
13891
13945
13730
13634
13709
13691
13666
13943
13873
13789
13782
13727
log10(TPM+1)

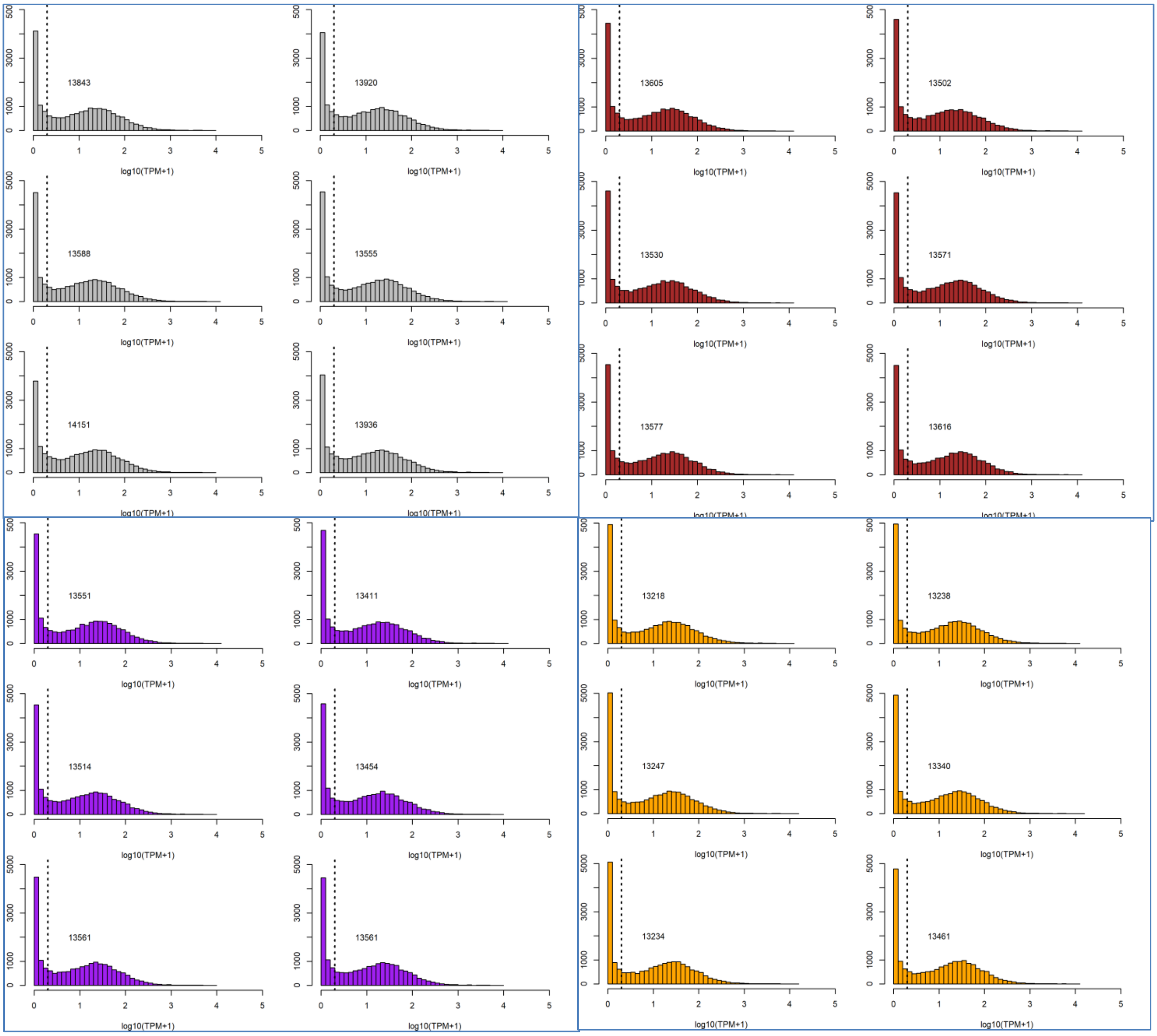


**Fig. S15. Distribution of transcripts expression.** For each condition, the TPM distribution in log10 scale is shown for the six corresponding samples (three independent transfection experiment, each performed in duplicates). Within each condition (*i.e.*, color), pairs of duplicates are displayed on the same row. The number of transcripts expressed above 1 TPM (dotted vertical line) is indicated. Colors are as follows: Shble#1 in pink, Shble#2 in green, Shble#3 in light blue, Shble#4 in red, Shble#5 in grey, Shble#6 in brown, EMPTY in purple and Mock in orange.

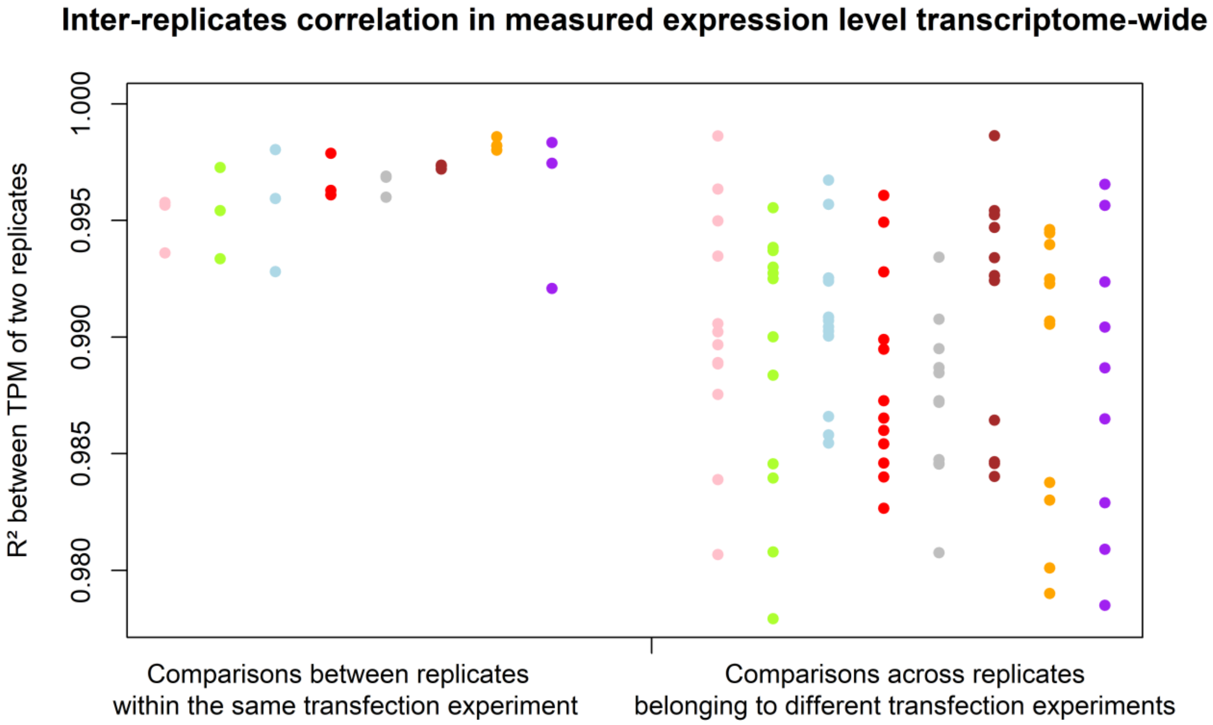


**Fig S16. Inter-replicates agreement in measured expression level transcriptome wide.** Three distinct transfection experiments were performed, each consisting of two duplicates per construct transfected. $R^2$ of pairwise comparisons between the two duplicates performed during the same transfection experiment (*left*) and between replicates belonging to different transfection experiments (*right*) were computed and compared. This was done to verify that it was possible to collapse measures obtained from duplicates of the same transfection experiment, thereby reducing the number of samples from 48 to 24. All data presented in the manuscript come from the 24 "collapsed" transcriptomic samples.

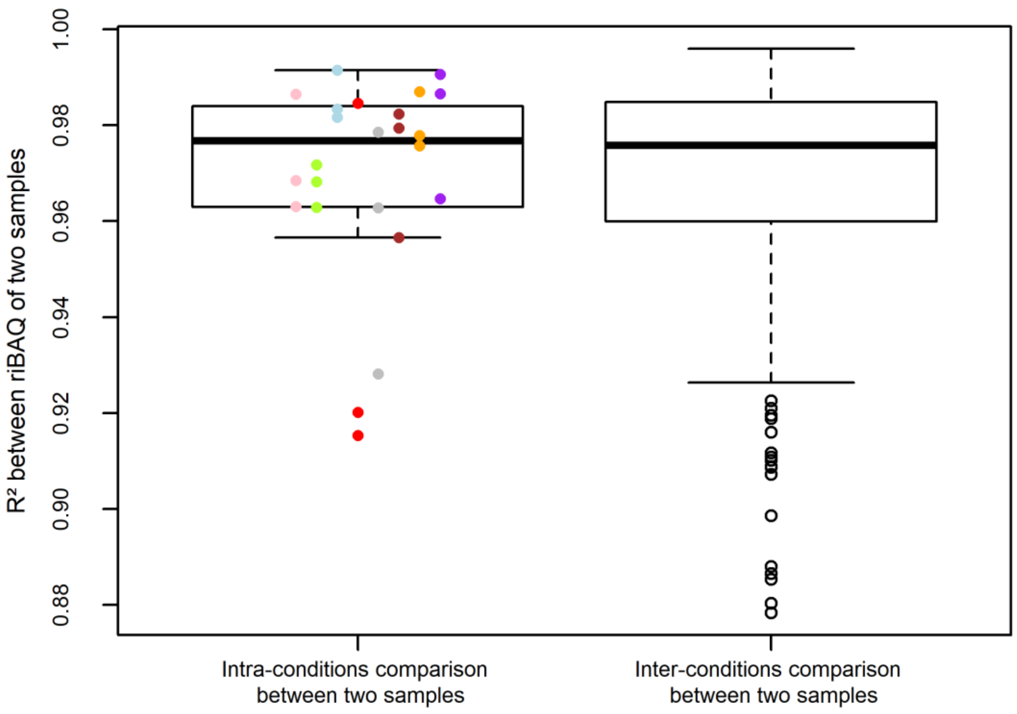


**Fig. S17. Inter-replicates agreement between proteomes.** Three independent transfection experiments followed by LC-MS/MS were performed for each of the eight conditions. $R^2$ of pairwise comparisons between proteomic expression patterns of samples from the same condition (*left;* condition-specific colors) or from different conditions (*right*) were computed. riBAQ values were used, with values corresponding to proteins not detected in one or more samples set to zero in these samples. Colors are as follows: Shble#1 in pink, Shble#2 in green, Shble#3 in light blue, Shble#4 in red, Shble#5 in grey, Shble#6 in brown, EMPTY in purple and Mock in orange.

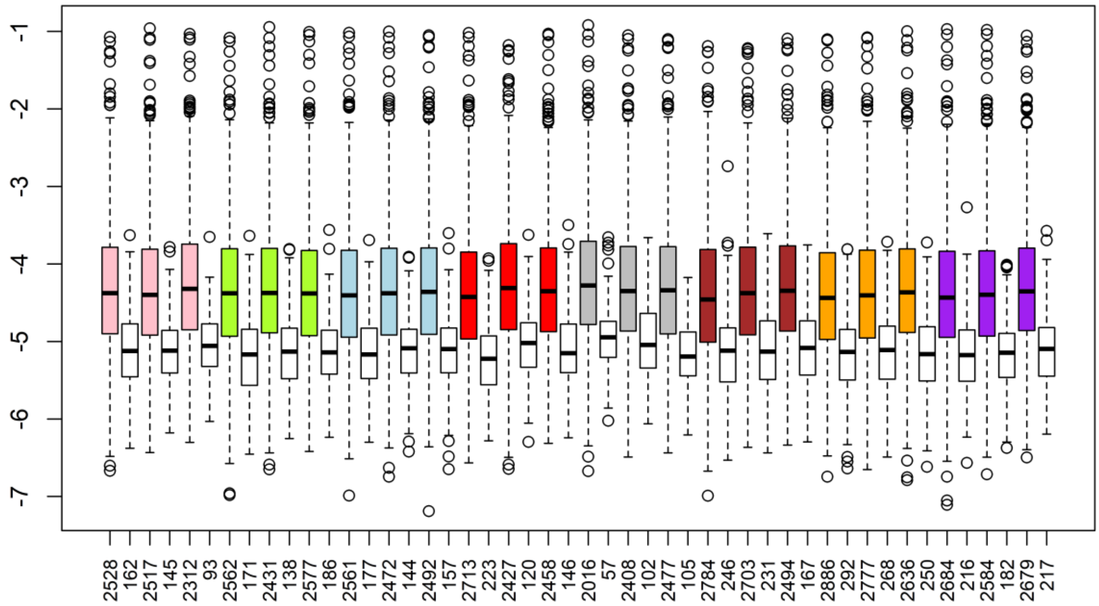


**Fig. S18. Number of detected proteins per sample and difference in expression between proteins not consistently detected in all samples of one condition (at least) and all proteins.** In this graph, boxplots are grouped by pair: colored boxplots represent expression of all detected protein in a given sample and white boxplots represent expression of all detected proteins in the same sample that belong to the set of 369 proteins displaying a qualitative pattern of expression according to the transfected construct (see the Methods section of the manuscript and Table S3). The number of genes included is added to the x-axis below each boxplot. The y-axis represents expression level in log10 riBAQ. The color code for the different conditions is as follows: Shble#1 in pink, Shble#2 in green, Shble#3 in light blue, Shble#4 in red, Shble#5 in grey, Shble#6 in brown, EMPTY in purple and Mock in orange.

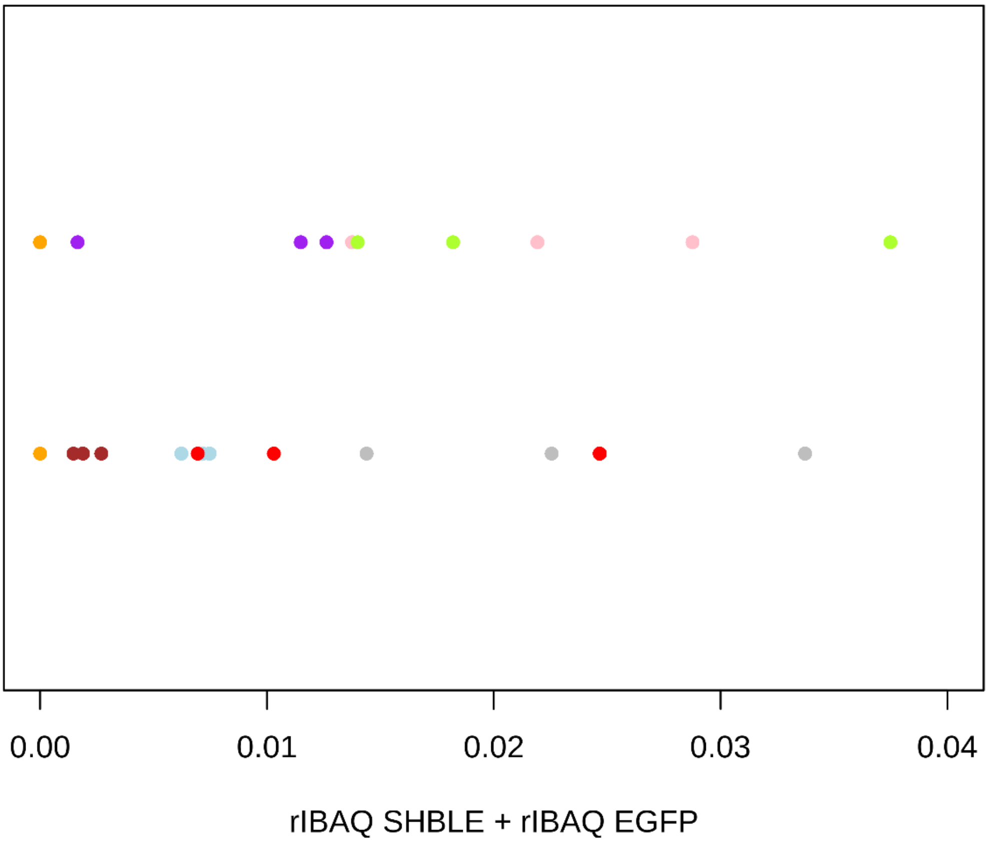


**Fig. S19. Sum of heterologous proteins expressed from samples transfected with over-humanized versions (upper row) or with non-humanized versions (lower row).** The sum of EGFP and SHBLE proteins (expressed as riBAQ values) by sample is reported. The two ranges of values shown served as the explanatory variable for regressions of riBAQ/TPM for indiviual genes versus total heterologous protein expression level (Figure 3 of the manuscript). Colors are as follows: Shble#1 in pink, Shble#2 in green, Shble#3 in light blue, Shble#4 in red, Shble#5 in grey, Shble#6 in brown, EMPTY in purple and Mock in orange. EMPTY samples (purple dots) were intentionnaly grouped in the upper row with Shble#1 (pink) and Shble#2 (green) samples because codon usage bias in *egfp* is also over-humanized.
